# Differential serum binding patterns predicting healthy subjects and cancer patients

**DOI:** 10.64898/2026.05.26.727832

**Authors:** Beatrice Cavalluzzo, Biancamaria Cembrola, Simona Mangano, Andre Belli, Francesco Izzo, Rosa D’Angelo, Maria Grazia Chiofalo, Cira Antonietta Forte, Alessandro Morabito, Alessandra Calabrese, Michelino De Laurentis, Vito Vanella, Paolo Ascierto, Fernanda Picozzi, Ottavia Clemente, Nicola Martucci, Ettore Pavone, Edoardo Mercadante, Franco Ionna, Maria Cristina Lucarelli, Angela Mauriello, Concetta Ragone, Limin Wang, Changzhi Ma, Yongmei Zhao, Xin Wei Wang, Maria Tagliamonte, Luigi Buonaguro

## Abstract

A viral exposure signature (VES) has been previously described predicting the development of Hepatocellular carcinoma (HCC) in at-risk patients. This has been achieved by a serological profiling of the viral infection history using a synthetic human virome including >100k epitopes (VirScan).

In the present study we applied the same VirScan strategy to identify a differential serum binding pattern (DSBP) for classifying patients of different cancer types from healthy individuals. In particular, the healthy group included both age-matched (ADULTS) as well as elderly (ELDERS) individuals, the latter counting also nonagenarians and centenarians.

The class comparison performed with serological data show DSBPs supporting class predictions, as confirmed by the receiver operating characteristic (ROC) curve analysis. Antibody responses supporting the class predictions are specific to peptides from persistent herpesviruses, acute-infecting viruses and, consistently in all comparisons, human respiratory syncytial virus (HRSV). Strikingly, the DSB of the ELDERS vs. CANCER comparison is characterized by higher titers in the healthy subjects; on the contrary, the DSB of the ADULTS vs. CANCER comparison is characterized by lower titers in the healthy subjects.

Overall, the results show a differential serological binding pattern predicting healthy individuals (ADULTS or ELDERS) from patients with different types of cancer. Such results provide the first evidence suggesting a close link between anti-microbial immunity and cancer development. They may be of the highest relevance in terms of predictive, diagnostic and/or prognostic impact in oncology.

## Introduction

The correlation between levels circulating immunoglobulins (Igs) and cancers has been previously evaluated with contrasting results suggesting either promoting or a protecting effect (1–4). Moreover, there are evidences that the density of the intra-tumoral IgG could be prognostic markers in patients with cancer (5,6). None of these studies, however, has explored the specificity of such antibodies. Alternatively, a correlation between Antibodies specific to cancer-associated agents (i.e. helicobacter pylori, hepatitis B virus, human papillomavirus) has been consistently reported (reviewed in 7).

The present study stems from the concept that, independent from any causative correlation, all viruses and bacteria are known to affect human health by altering host immunity. Consequently, not only the levels but, more importantly, the antigen specificity of Igs may play a crucial role in the pathogenesis of human chronic diseases, including cancer (8–10). Indeed, several pathogenic and non-pathogenic microbes may shape host immunity, altering its response to new infections and cancer risk. Consequently, microbes, even when cleared, leave unique immunological footprints that can affect host susceptibility to cancer. Such a distinctive signature may represent either an unfavorable or a protective prediction pattern.

A viral exposure signature (VES) has been previously described to be associated with Hepatocellular carcinoma (HCC), discriminating patients from at-risk or healthy volunteers. The synthetic virome technology, VirScan, based on a high-throughput sequencing method, was used to detect the exposure history to all known human viruses (11). The signature identified cancer patients prior to a clinical diagnosis and was superior to alpha-fetoprotein (12).

Alternatively, we have proposed that immune responses elicited by microbial antigens (MoAs), mimicking tumor antigens (TAAs), may protect from cancer development or rapid progression (13). Several recent reports have shown microorganism-associated antigens (MoAs) mimicking tumor-associated antigens (TAAs) (molecular mimicry) along with induction of cross-reacting T cells with cytotoxicity activity (14–25).

More recently, we have also shown that the SARS-CoV-2 preventive vaccine, besides the humoral response, is able to elicit also a T cell response which cross-react with homologous TAAs (26). This would suggest a “preventive” anti-cancer effect of the immune response induced by the global vaccination during the COVID-19 pandemic.

In this framework, we hypothesize that a unique broad anti-microbe immunity may protect individuals during their life from cancer development. To this aim, a cohort of cancer patients with different diagnosis was compared to a cohort of healthy subjects adopting the VirScan synthetic virome technology. In the latter cohort, nonagenarians and centenarians were also included as “true cancer-free” subjects, assuming that age-matched healthy subjects may only be apparently or momentarily cancer-free.

Such an agnostic approach has identified a differential viral exposure signature (VES), deriving from virus-host interactions, which may represent a key advancement in the definition of an immunological biomarker of protection from cancer as well as in the knowledge of potentially modifiable factors.

## Methods

### Patient Cohorts

The study enrolled 359 participants: 187 healthy controls (HEALTHY cohort) and 172 cancer patients (CANCER cohort). The healthy cohort included two age-based subgroups: 82 adults <70 years old (ADULTS subgroup) and 106 elders >80 years old, including nonagenarians and centenarians (ELDERS subgroup). The participants in the ADULTS and the CANCER cohorts were enrolled at the National Cancer Institute “Pascale” in Naples, ITALY. The participants in the ELDERS cohort were enrolled in 14 villages of the “Comunità Montanta del Fortore”, Benevento – IT.

### Sample collection

Peripheral blood was obtained by venipuncture from each individual upon signing informed consent. The blood was diluited 1:2 with DPBS 1X (Dulbecco’s Phosphate buffered saline 1X) and centrifuged at 1200 g for 10 minutes at room temperature. Plasma in the upper phase was collected, aliquoted and stored at -80°C until use.

### Samples’ Preparation for VirScan analysis

Human plasma samples were completely thawed and mixed thoroughly before aliquoting to ensure IgG homogeneity and dissolve any precipitate formed during storage. For each sample, 20 µL was transferred into round-bottom (V-shape) 96-well PCR plates (Bio-Rad, Cat #HSP9601). Samples were loaded sequentially by row (A1 to A12, B1 to B12, etc.). Each full plate contained 87 samples (wells A1 to H3), leaving wells H4 to H12 empty for downstream technical controls. Plates were sealed at room temperature with an adhesive film applicator and specific sealing sheets, then immediately frozen and shipped on dry ice.

### Pan-microbial serological profiling of serum samples

Serum samples were profiled by phage immunoprecipitation sequencing. The phage library used for immunoprecipitation includes 115,753 synthetic peptides from known human viruses, fungi, and bacteria genomes. A total of 2 microgram immunoglobulins from each sample were mixed with the 1 milliliter phage library and 40 microliters of magnetic Dynabeads, with technical duplicates. The bound complexes were then washed and recovered in 40 microliters of water and boiled for sequencing library construction. Sequencing libraries were constructed by two rounds of PCR and necessary DNA purification steps. The DNA libraries were then pooled and sequenced on an Illumina NovaSeq platform.

### Bioinformatic processing of sequencing data

Raw sequencing data was de-multiplexed and mapped to the reference genome using Bowtie 2. Zero-inflated p-values of each epitope were calculated based on the read counts of technical duplicates. The binding scores of each virus strain and species were then accounted using p-value data. Read counts of each epitope were also used to calculate the Epitope Binding Signal (EBS) of each epitope and the data was further filtered with p-value data and removed the cross-reactive epitopes.

### Serological Profiling and Data Analysis

We performed high-throughput serological profiling of viral infection histories using the VirScan platform. This system screens antibody binding against a synthetic human virome of 82,134 unique peptides from 1,558 organisms and 444 species (including 241 viral and 203 bacterial/parasitic species). The resulting single-channel raw intensity screening data, representing antibody binding titers for each peptide, were processed with BRB-ArrayTools (v4.6). To allow mathematical operations within the data matrix, zeros were replaced with a baseline threshold of 0.1 before formal collation. Unique peptide identifiers were aligned horizontally alongside the serological intensity profiles. We bypassed data normalization and intensity filtering during initial collation to preserve the full spectrum of raw serological responses. Class Comparison analysis was performed across cohorts defining the phenotype classes in the experiment descriptors. Class prediction analysis was performed building predictive models using the identified DSBPs to classify patients versus healthy controls. Model accuracy and robustness were validated through a 10-fold cross-validation scheme. We evaluated classification sensitivity and specificity by calculating the Area Under the Curve (AUC) via Receiver Operating Characteristic (ROC) curve analysis. For all analyses the threshold p-value was set at <0.01.

## Results

### Cohort study

187 healthy subjects and 172 cancer patients were enrolled for the study. The age distribution in the two cohort groups showed two different patterns. The cancer cohort showed a unimodal distribution with a median at 64.07 years, while the healthy cohort showed a bimodal distribution with an overall median at 71.74 years (p-value <0.001) (Fig. 1A). The latter bimodal distribution was the result of the experimental design. Indeed, the two peaks showed a median at 50.3 and at 88.1 years (Fig. 1B), corresponding to the two sub-cohorts of volunteers selected for the present study and defined as ADULTS (<70 years old) and ELDERS (>80 years old) (p-value <0.0001).

**Fig. 1.**
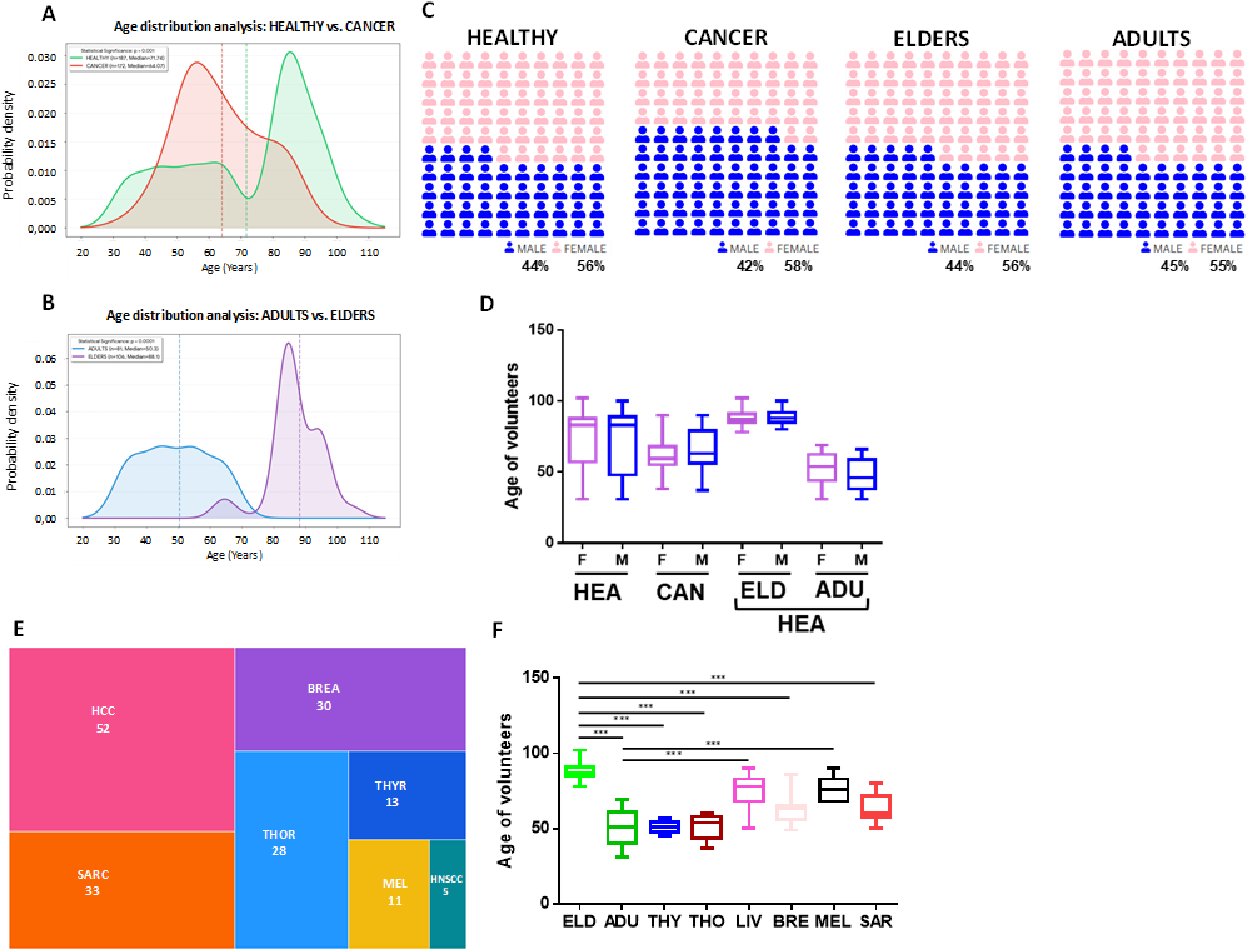
Cohort groups in the study. Age distribution in both HEALTHY and CANCER subjects (A) and within the HEALTHY subjects (ELDERS and ADULTS) (B). The male to female distribution is represented in panel C and D. The number of patients from each cancer type and their age distribution is represented in panel E and F.

In the HEALTHY and CANCER cohort groups, the percentage of males and females was almost perfectly mirrored, with males prevalent in the CANCER group (58%) and females prevalent in the HEALTHY group (56%). Within the latter group, females were prevalent in both ELDERS and ADULTS (55-56%). Overall, the average age within each group showed a very good balance between males and females (Fig. 1C - D).

The CANCER group included patients from 6 histotypes and the numerosity of enrolled volunteers ranged from 52 with hepatocellular carcinoma (HCC) to 11 with melanoma (MEL). The average age of the cancer patients showed three groups: 1) thyroid and thoracic cancers (50.8 years); 2) breast cancer and sarcoma (62.8 years); and 3) HCC and MEL (75.9 years) (Fig. 1E - F) (Table 1).

**Table 1.** Clinical characteristics of the Subjects.

| Variable | ELDERS<br>(n = 106) | ADULTS<br>(n = 81) | THYR<br>(n = 13) | THOR<br>(n = 29) | SARC<br>(n = 33) | BREA<br>(n = 30) | HCC<br>(n = 51) | MEL<br>(n = 11) | HNSCC<br>(n = 5) |
| --- | --- | --- | --- | --- | --- | --- | --- | --- | --- |
| **Age - year** |  |  |  |  |  |  |  |  |  |
| **Median**<br>**(range)** | 87<br>(78 – 102) | 51<br>(31 – 69) | 51<br>(45 – 57) | 54<br>(38 – 60) | 60<br>(50 – 80) | 63<br>(50 – 79) | 78<br>(50 – 90) | 76<br>(68 – 90) | 57<br>(55-60) |
| **p-value** |  |  |  |  |  |  |  |  |  |
| **vs. ELDERS** | 1 | <0.0001 | <0.0001 | <0.0001 | <0.0001 | <0.0001 | <0.0001 | <0.001 | <0.0001 |
| **vs. ADULTS** | <0.0001 | 1 | 0.94 | 0.97 | <0.0001 | <0.0001 | <0.0001 | <0.0001 | <0.0001 |
| **Sex – no. (%)** |  |  |  |  |  |  |  |  |  |
| **Female** | 59 (55.7%) | 45 (55.6%) | 6 (46.1%) | 15 (51.7%) | 19 (57.6%) | 30 (100%) | 10 (19.6%) | 6 (54.5%) | 2 (40%) |
| **Male** | 47 (44.3%) | 36 (44.4%) | 7 (53.8%) | 14 (48.3%) | 14 (42.2%) |  | 41 (80.4%) | 5 (45.5%) | 3 (60%) |
| **p-value** |  |  |  |  |  |  |  |  |  |
| **vs. ELDERS** | 1 | 0.97 | 0.95 | 0.97 | 0.96 |  | <0.0001 | 0.99 | 0.87 |
| **vs. ADULTS** | 0.97 | 1 | 0.96 | 0.98 | 0.97 |  | <0.0001 | 0.99 | 0.82 |

### Serological analysis by VIRSCAN

Serum from each individual was screened against 82,134 peptides derived from 1558 organisms and 444 species, including 241 viral and 203 bacterial/parasite species.

Sera from both HEALTHY and CANCER groups showed reactivities toward only 11.55% peptides with a very broad percentage of responsive subjects. Considering a threshold of reactivity in >50% of subjects in both groups, the reacting peptides are 43 (HEALTHY) and 42 (CANCER) and derive from 8 pathogens with a very similar distribution. Three are persistent herpesviruses (HHV-1, HHV-5 and HHV-4, highest-to-lowest), four acute-infection viruses (Rhinovirus A, HRSV, Rhinovirus B and Adenovirus C, highest-to-lowest) and one bacterium (*Streptococcus Pneumoniae*) (Fig. 2 A and B). Strikingly, the top 10 peptides with the highest responses derive from *Streptococcus Pneumoniae* (5 in HEALTHY; 4 in CANCER), HHV-1 (2 in HEALTHY; 3 in CANCER), HHV-4 (1 in HEALTHY; 1 in CANCER), HHV-5 (2 in CANCER) and Rhinovirus A (2 in HEALTHY) (Fig. 2 E and F). Within the HEALTHY group, the ELDERS and ADULTS subgroups show a significantly different picture. The reacting peptides are 54 (ELDERS) and 34 (ADULTS), but the distribution of the 8 pathogens is greatly different (Fig. 2 C and D) with a very limited representation of the HSRV and higher representation of Rhinovirus A, combined to a lower representation of HHV-1, in ADULTS. Such a difference is reflected also in the top 10 peptides with the highest responses. Indeed, besides the same top 2 peptides from *Streptococcus Pneumoniae* in both groups, the remaining 8 peptides are totally different with the single exception of a peptide from HHV-4. The ELDERS respond to HHV-1 (5 peptides) and HHV-5 (2 peptides); the ADULTS respond to *Streptococcus Pneumoniae* (4 peptides), Rhinovirus A (2 peptides) and Rhinovirus B (1 peptide) (Fig. 2 G and H).

**Fig. 2.**
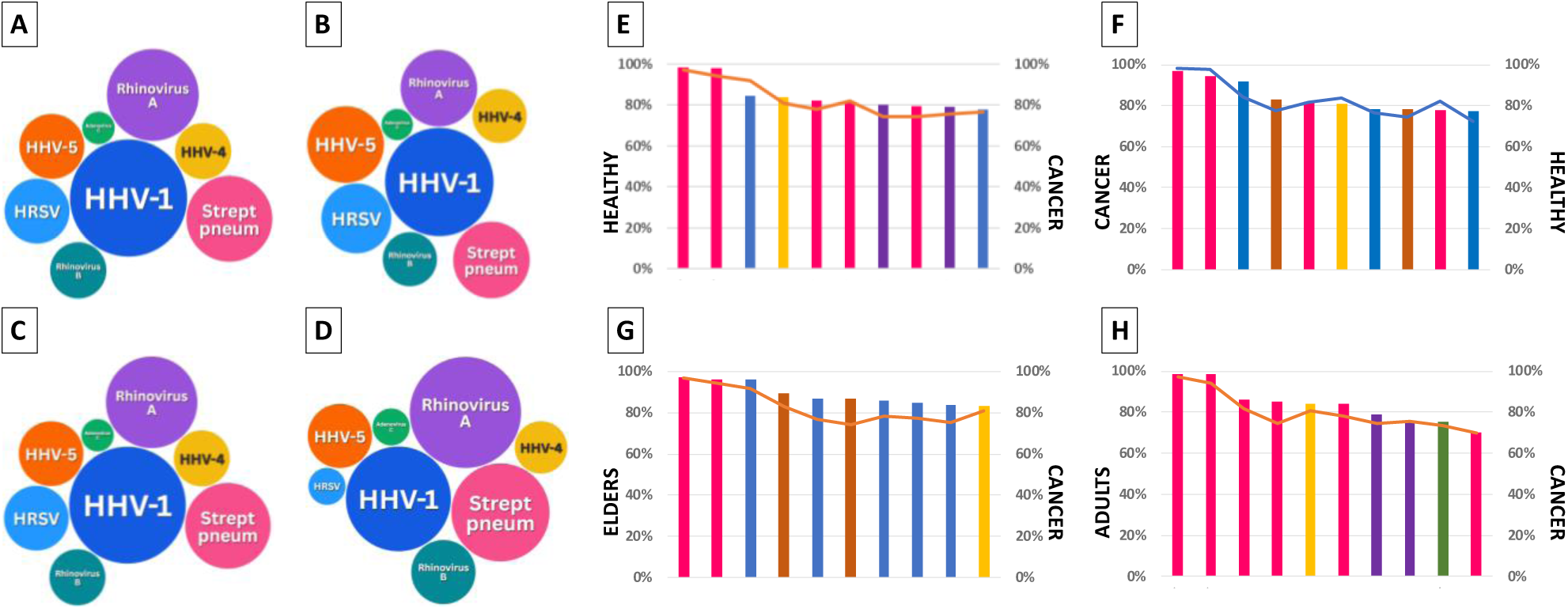
Prevalent serum binding in the enrolled subjects. Numerosity of microbes from which are derived the peptides with >50% of responders in HEALTHY (A) and CANCER (B) subjects; as well as in the subgroups of ELDERS (C) and ADULTS (D). The top 10 peptides with responders in HEALTHY (E) and CANCER (F) subjects; as well as in the subgroups of ELDERS (G) and ADULTS (H). The same color code is applied to panels A-D and E-H.

### Differential serum reactivity in the two groups

In order to find a differential pattern of serum reactivity in the groups, a class comparison was performed with serological data obtained in all 172 CANCER subjects vs. either all 187 HEALTHY or the 105 ELDERS and the 82 ADULTS, separately. The analysis comparing the entire number of subjects in the two groups identified a set of 15 peptides differentially recognized by sera (p-value <0.01), of these only 5 show a significant down (nr. 4) or up (nr.1) Log2 fold change in HEALTHY subjects (p-value 0.001) (Fig. 3 A_1_ – A_2_). However, such differential serum binding (DSB) is not sufficient to support a class prediction, as shown by the receiver operating characteristic (ROC) curve analysis (Fig. 3 A_3_).

**Fig. 3.**
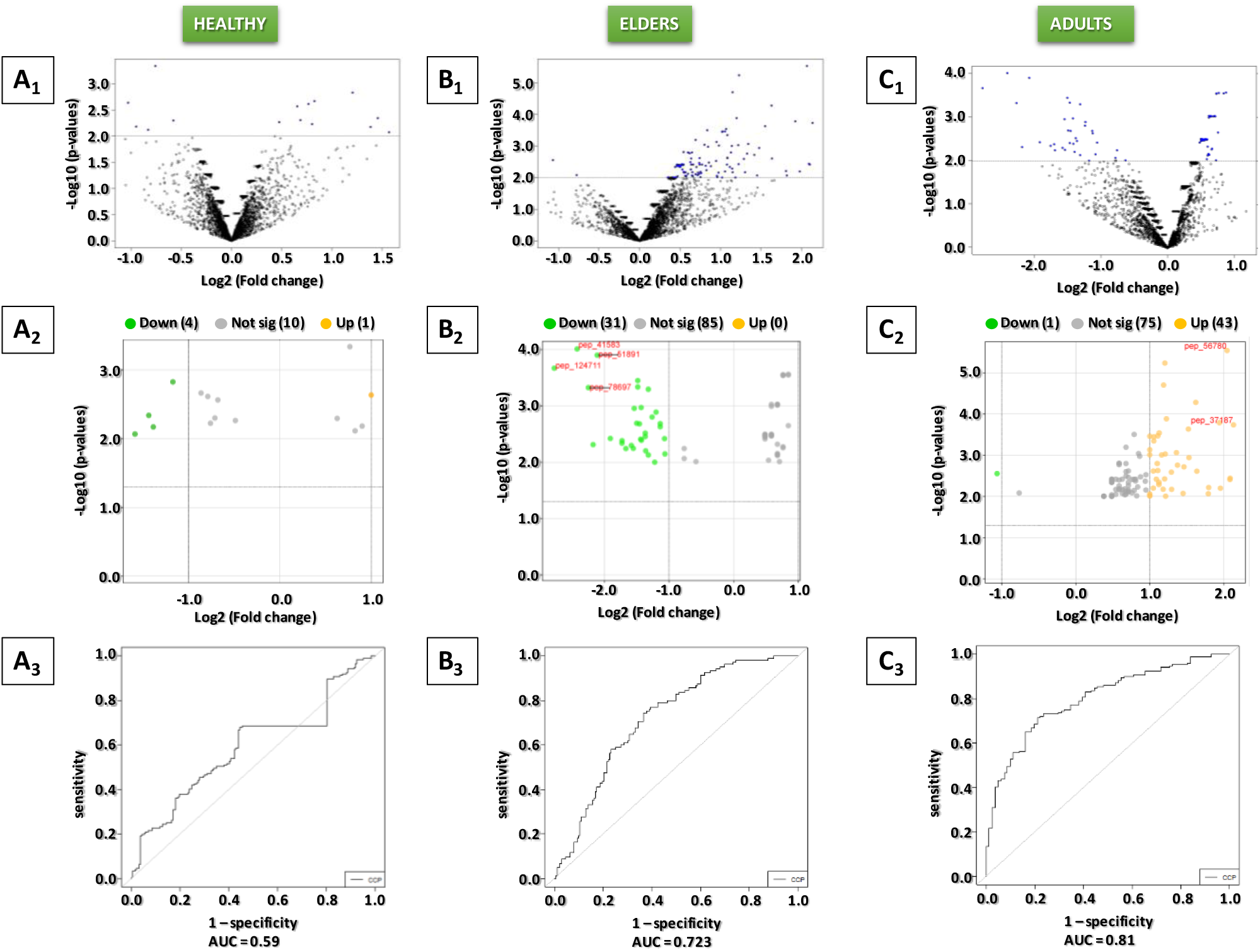
Differential serum binding pattern (DSBP) with all CANCER patients. Class comparison analysis based on antibody titers specific for the VirScan peptide array. All HEALTHY subjects vs. all CANCER patients (A_1_ - A_2_); ELDERS vs. all CANCER (B_1_ - B_2_); ADULTS vs. all CANCER (C_1_ - C_2_). The ROC curves derived from class prediction analysis for each of the comparisons are shown (A_3_ – C_3_). AUC = area under the curve.

On the contrary, the same analysis performed comparing the CANCER subjects with the ELDERS or the ADULTS, independently, show a completely different outcome. Indeed, 44 and 31 peptides differentially recognized by sera are identified in ELDERS and in ADULTS, respectively (p-value <0.01) (Fig. 3 B_1_ and C_1_). However, compared to CANCER subjects, the ELDERS show a higher Ab response to 43/44 peptides, while the ADULTS show always a lower titer (Fig. 3 B_2_ and C_2_). Irrespective of such a symmetrical result, the differential serum binding pattern (DSBP) supports a class prediction in both comparisons (ELDERS vs. CANCER; ADULTS vs. CANCER), as shown by the receiver operating characteristic (ROC) curve analysis (Fig. 3 B_3_ and C_3_).

The peptides identified in the DSBP and supporting the class prediction in both comparisons derive from 27 microorganisms (24 viruses, 2 bacteria, 1 protozoan), of which seven viruses are in common. The largest number of peptides in both comparisons are from the Human respiratory syncytial virus (HRSV), while the second largest number of peptides is from the Human herpesvirus 5 (ELDERS vs. CANCER) and from the Influenza A virus (ADULTS vs. CANCER). Overall, the peptides from the persistent herpesviruses are predominant in the ELDERS (20/43) compared to the ADULTS (5/31) (Fig. 4).

**Fig. 4.**
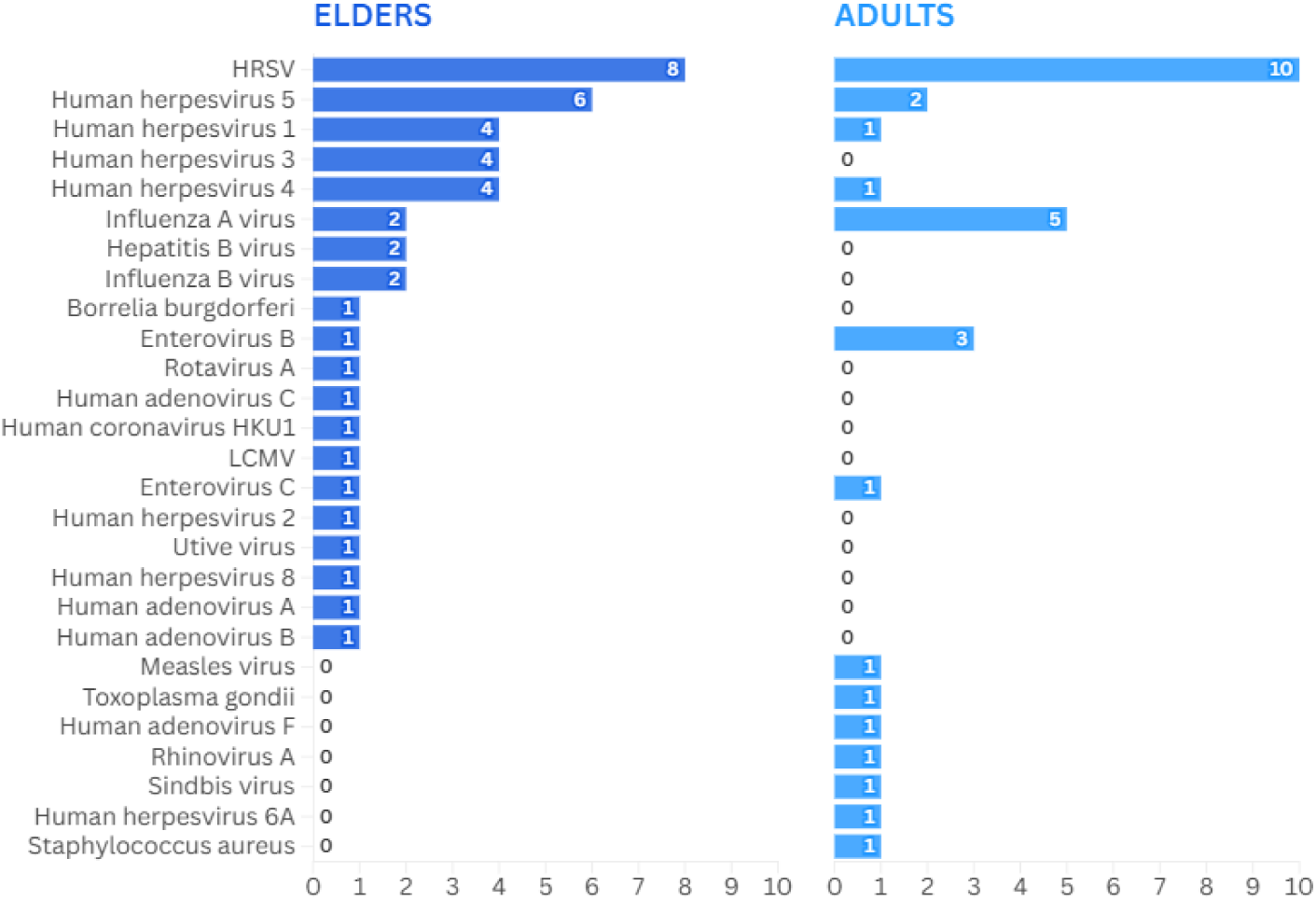
Peptides identified in the DSBP with all CANCER patients. Numerosity of peptides derived from the indicated microbes found in the DSBP of the comparisons ELDERSvs.CANCER as well as ADULTSvs.CANCER.

The percentage of subjects reacting to each of such peptides provides a picture of the immunological relevance. In the ELDERS vs. CANCER comparison, although the Log2fc of the Ab responses are >1 and supported by a significant p-value, none of the peptides is recognized by a relevant percentage of HEALTHY subjects (e.g. >75%) in one group and consistently in a much lower percentage (e.g. <30%) in patients from all cancers. Indeed, the peptide 30608 (Human herpesvirus 1) is the only one recognized by 77.36 of healthy subjects and by an average of 46.1% of cancer patients (30.3% SARC - 60.8% HCC). Very interestingly, quite few peptides are not recognized by patients from one or more cancer types. Melanoma patients do not have antibodies to 26/44 peptides and 13 of these are from herpesviruses; moreover, 21 of the 26 peptides are not recognized also by patients with one or more (up to four) other histotypes. Indeed, the peptides 65050 (HHV-8), 9115 (Adeno A) and 8005 (Adeno B) are not recognized by patients with 4 histotypes in different combinations, namely MEL, BRE, HCC, SARC and THYR patients (Suppl. Table 1).

In the ADULTS vs. CANCER comparison, the picture is quite similar and none of the peptides is recognized by a relevant percentage of subjects (e.g. >75%) in one of the groups and consistently in a much lower percentage (e.g. <30%) of subjects in the compared one. Also in this comparison, a number of peptides (11/31 peptides) are not recognized by melanoma patients and 6 of these 11 peptides are not recognized by patients with one or more (up to three) other histotypes. The peptides 82438 (Measle virus), 25542 (HHV-1) and 94230 (Influenza A) are not recognized by patients with 3 histotypes in different combinations, namely MEL, BRE, HCC, and THYR patients (Suppl. Table 1). However, in both ELDERS vs. CANCER and ADULTS vs. CANCER combinations the responses to the same peptides is observed in <10% of the HEALTHY subjects.

### Differential serum reactivity in healthy subjects and patients with different cancer histotypes

The subsequent class comparison analyses were performed to identify a differential pattern of serum reactivity in all 187 HEALTHY or the 105 ELDERS and the 82 ADULTS, respectively, vs. patients with the different histotypes. Each one-to-one comparison identified a set of peptides differentially recognized by sera (p-value <0.01). Interestingly, in ELDERS subjects the DSBP was characterized by higher Ab responses when compared to breast ca, liver ca and sarcoma patients as well as lower Ab responses when compared to melanoma, thyroid ca and thoracic ca patients. Conversely, in ADULTS subjects the DSBP was characterized by lower Ab responses when compared to each group of cancer patients (Suppl. Fig 1). However, such DSBP support a class prediction with an area under the curve (AUC) ≥70% only in the comparisons ELDERS vs. HCC, ADULTS vs. THYROID, EDELRS/ADULTS vs. SARCOMA and EDELRS/ADULTS vs. THORACIS as shown by the ROC curve analysis (Suppl. Fig 2).

### DSBP in ELDERS vs. HCC comparison

The DSBP supporting the class prediction in the comparison ELDERS vs. HCC is based on 47 peptides. Of these, 40 show a serological binding titer higher in ELDERS and 7 higher in HCC (Fig. 5 A_1_ – A_3_; Suppl. Table 2). Overall, the peptides derive from 25 microorganisms (23 viruses, 2 bacteria) and the top three are HRSV (6 peptides), HHV-4 (5 peptides) HHV-1 (4 peptides) (Fig. 5B). The median percentage of ELDERS reacting to peptides from each microorganism is very variable, ranging from 66.5% (HHV-1) to 1.4% (Enterovirus A); in parallel, the data from the HCC patients show significantly lower median values for most of the microorganisms, except for HHV-1, HHV-4, Rhinovirus A and Adenovirus C (Fig. 5C). Indeed, in all the latter cases, the median of percentages of subjects reacting to the peptides are not statistically different from the ELDERS group. Regardless the statistics, the Log2 fold change in the Ab titers reacting with the individual peptides from each microorganism, is highly relevant between subjects of the two groups. In particular, with the only exception of the peptides from the Enterovirus A, in all other cases the fold change is higher in ELDERS than in HCC subjects (Fig. 5D). The two peptides recognized by >75% of ELDERS derive from HHV-1 (pep_80349 and pep_30608) and, although a similar percentage of responders are observed also in HCC patients, the Log2 fold change in the Ab titers is >2.6. Strikingly, in all of the HCC patients the serum reactivity to some peptides is lacking, namely pep_83385 (Influenza B), pep_120234 (Borrellia burgdorferi), pep_111489 (Utive Virus), pep_9115 (Adenovirus A) and pep_54814 (Papiine herpesvirus 2). The percentage of responders in the ELDERS is 11.5%, on average (8.49% - 14.15%) (Suppl. Table 2).

**Fig. 5.**
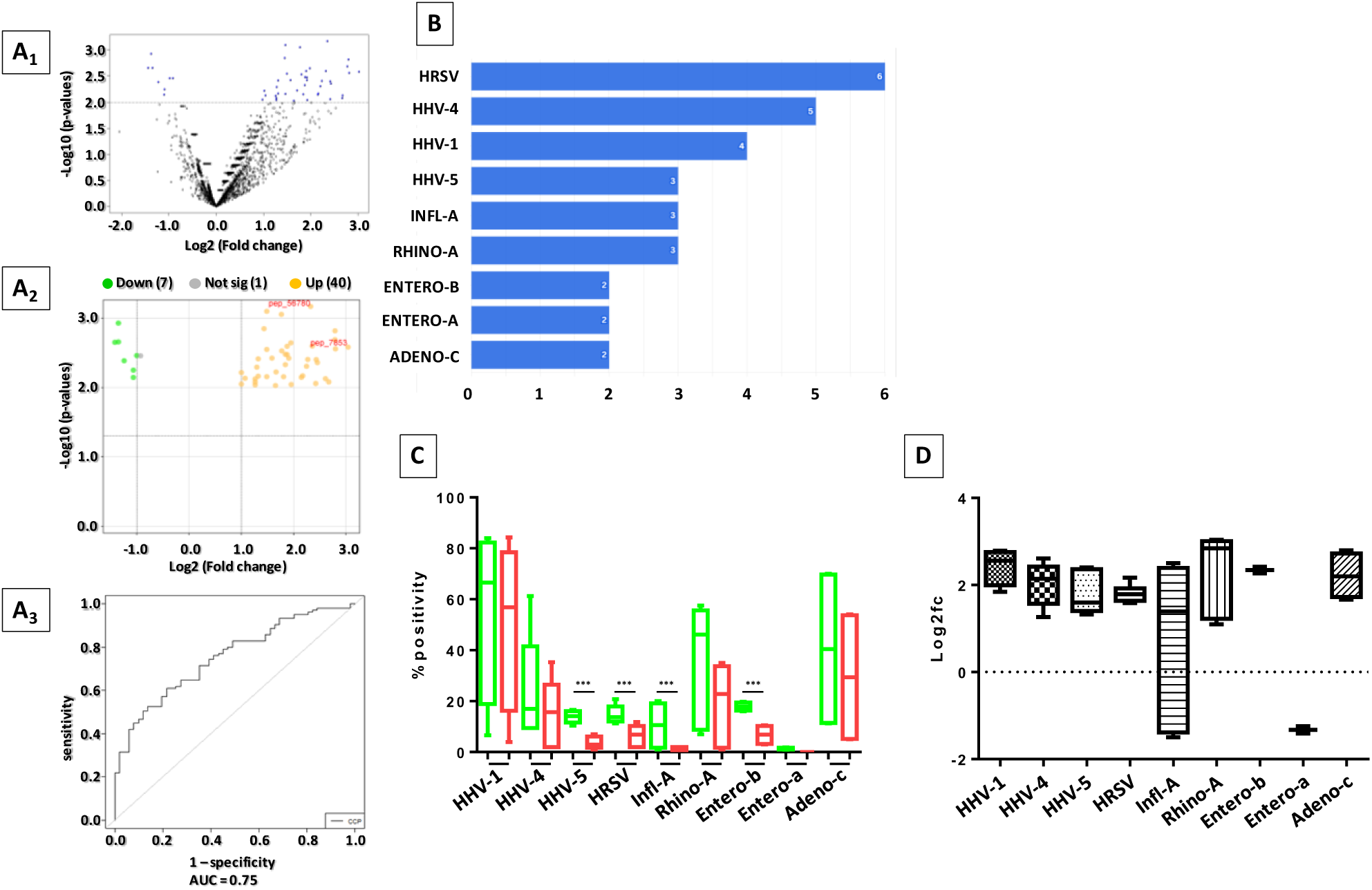
Differential serum binding pattern (DSBP) in ELDERSvs.HCC comparison. Class comparison and prediction analysis based on antibody titers specific for the VirScan peptide array in ELDERS vs. HCC patients (A_1_ – A_3_). Numerosity of peptides derived from the indicated microbes found in the DSBP (threshold of 2 peptides) (B). Percentage of responders in the two groups to all peptides from each microbial specie (C); differential binding titers between ELDERS vs. HCC patients (D).

### DSBP in ADULTS vs. THY comparison

The DSBP supporting the class prediction in the comparison ADULTS vs. THY is based on 145 peptides, all showing a serological binding titer lower in ADULTS than in thyroid cancer patients (Fig. 6 A_1_ – A_3_; Suppl. Table 3). Overall, they derive from 49 microorganisms (44 viruses) and the top three are HRSV (11 peptides), HHV-4 (10 peptides) and HHV-1 (9 peptides) (Fig. 6 B). The median percentage of ADULTS subjects reacting to peptides from each microorganism is very variable but always <20%, while the data from the THYR patients show significantly higher median values for all the microorganisms (Fig. 6 C). The Log2 fold change in the Ab titers reacting with the individual peptides from each microorganism is significantly lower in ADULTS than in THYR patients, particularly for the peptides from the Enterovirus B (average -4.9 Log2 fold change) (Fig. 6 D). None of the peptides is recognized by >75% of subjects, but top three peptides are from HHV-1 (the overlapping pep_24523 and pep_70404) and Staphylococcus aureus (pep_124711). In particular, the latter peptides is recognized by about 70% of thyroid patients and 29.3% of ADULTS (2.33 fold) (Suppl. Table 3). Strikingly, 30 peptides are not recognized by ADULTS and 16 of them are derived from Influenza A; notably, five or them are recognized by >20% of thyroid patients, 2 peptides from Influenza A, 1 from HBV, 1 from HHV-6B and 1 from Molluscum Contagiosum (Suppl. Table 3). On the contrary, 22 peptides are not recognized by thyroid patients and, also in this case, 13 of them are derived from Influenza A, with no overlapping with those not recognized by ADULTS. None of such peptides is recognized by >6.5% of ADULTS. Interstingly, 18 peptides are bound by a percentage of THYR patients ranging from 20 to 30% and a percentage <5% of ADULTS and 12 are from Influenza A (Suppl. Table 3).

**Fig. 6.**
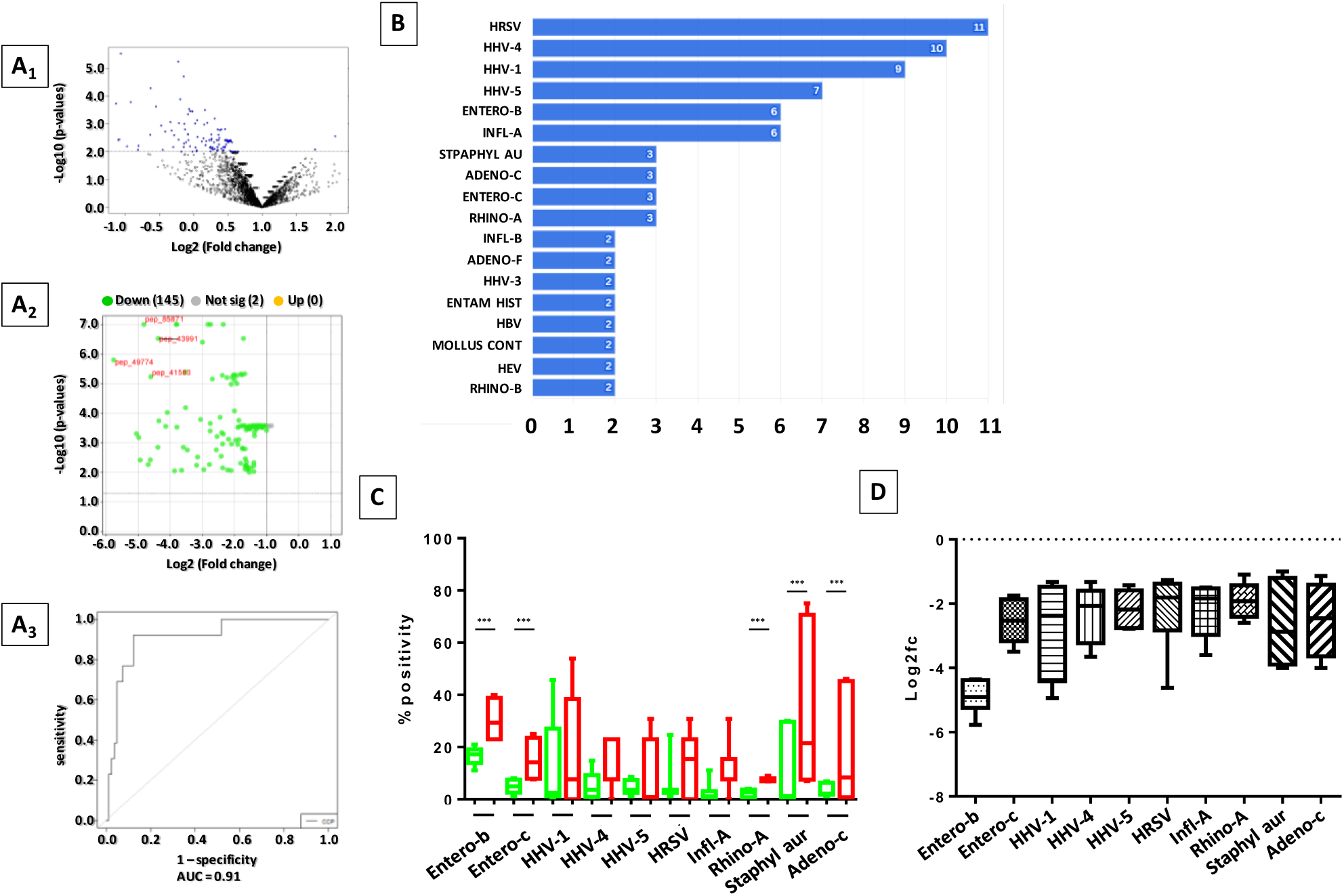
Differential serum binding pattern (DSBP) in ADULTSvs.THYR comparison. Class comparison and prediction analysis based on antibody titers specific for the VirScan peptide array in ADULTS vs. THYR patients (A_1_ – A_3_). Numerosity of peptides derived from the indicated microbes found in the DSBP (threshold of 2 peptides) (B). Percentage of responders in the two groups to all peptides from each microbial specie (C); differential binding titers between ADULTS vs. THYR patients (D)

### DSBP in HEALTHY (ELDERS and ADULTS) vs. THOR comparison

The comparisons between HEALTHY, ELDERS or ADULTS, and thoracic cancer patients (THOR) generated a DSBP supporting the class prediction in both analyses (ELDERS vs. THOR & ADULTS vs. THOR) and, in both cases, this is based on serological binding titers lower in the HEALTHY subjects than in the thoracic cancer patients (Fig. 7 A_1_ – A_3_; B_1_ – B_3_; Suppl. Table 4 and 5).

**Fig. 7.**
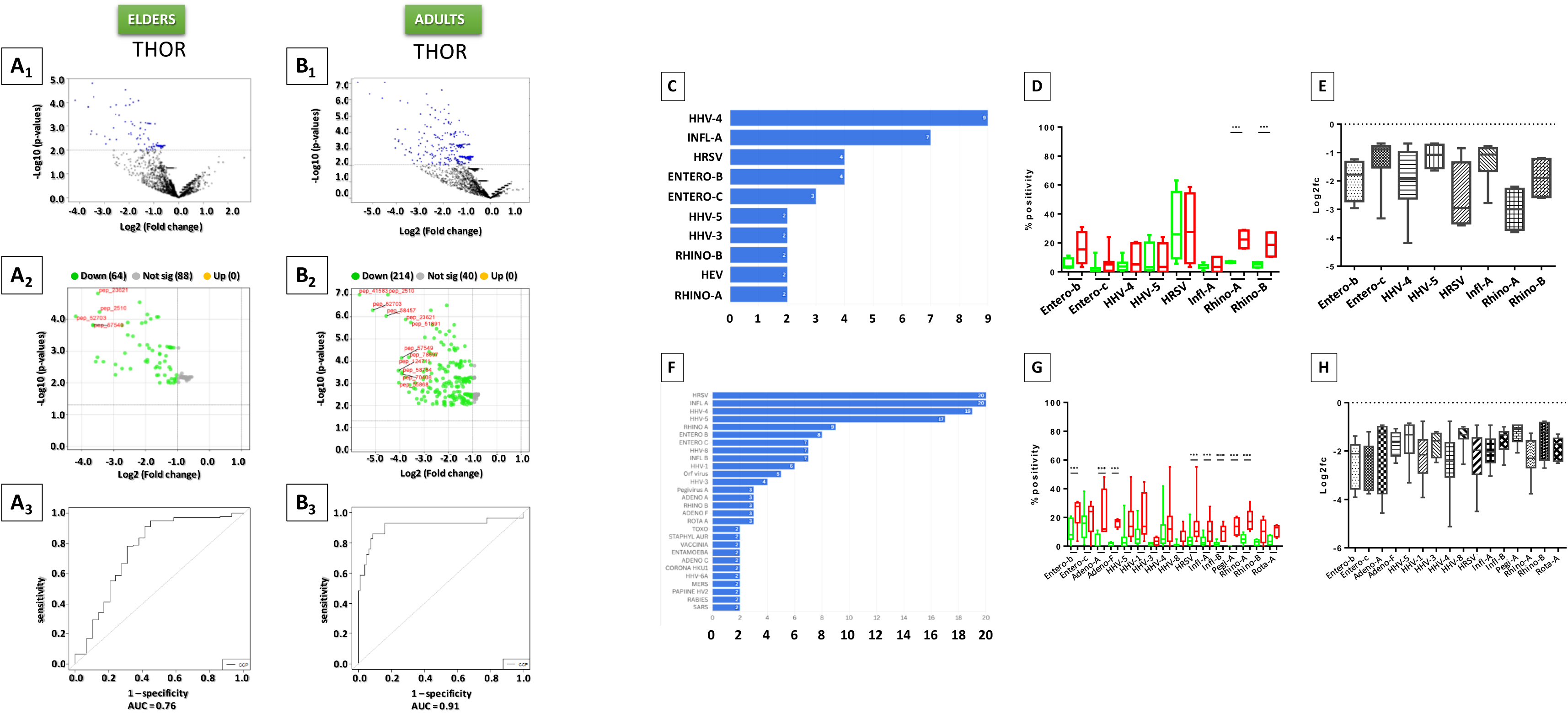
Differential serum binding pattern (DSBP) in HEALTHYvs.THOR comparison (C-H). Class comparison and prediction analysis based on antibody titers specific for the VirScan peptide array in ELDERS (A_1_ – A_3_) or ADULTS (B_1_ – B_3_) vs. THOR patients. Numerosity of peptides derived from the indicated microbes found in the DSBP (threshold of 2 peptides) (C and F). Percentage of responders in the two groups to all peptides from each microbial specie (D and G); differential binding titers between ELDERS (E) or ADULTS (H) vs. THYR patients.

The ELDERS vs. THOR DSBP is based on 64 peptides deriving from 31 microorganisms (30 viruses) and the top three are HHV-4 (9 peptides), Influenza A (7 peptides) and HRSV & Enterovirus B (4 peptides) (Fig. 7C). The median percentage of ELDERS subjects reacting to peptides from each microorganism is <10%, with only exception of HRSV (25.7%), while the data from the THOR patients show significantly higher median values for all the microorganisms. In particular, the median percentage of THOR responders to peptides from Rhinovirus A and B is significantly higher than the ELDERS (22.4% vs. 6.6% and 18.9% vs. 4.72%, respectively) (Fig. 7D). The median Log2fc change in the Ab titers reacting with the individual peptides from each microorganism is significantly lower in ELDERS than in THOR patients, particularly for the peptides from HRSV and Enterovirus B (Fig. 7 E). A single peptide from HRSV (pep_2496) is recognized by about 60% of subjects in both groups with a relevant lower Ab titer in ELDERS (Log2fc -3.56). Overall, the average percentage of responders in both groups is relatively low (6.48% ELDERS; 10.92% THOR patients). Only 5 peptides are recognized by >20% of ELDERS (3 HRSV, 1 Adenovirus A, 1 HHV-5) and by a similar or higher percentage of THOR patients; on the contrary, 12 peptides are recognized by >20% of THOR patients and 4 of them are recognized by <7% of ELDERS (2 HHV-4, 1 Rhinovirus A, 1 Rhinovirus B). A single peptide is not recognized by any subject in the ELDERS group (pep_125342 from Streptococcus pneumoniae) while 15 peptides are not recognized by any subject in the THOR group, but in both cases the percentage of responders in the other group is very limited (<5%) (Suppl. Table 4).

The ADULTS vs. THOR DSB pattern is based on 214 peptides deriving from 63 microorganisms (58 viruses) and, also in such comparison, the top three are HRSV and Influenza A (20 peptides), HHV-4 (19 peptides), HHV-5 (17 peptides) (Fig. 7F). The median percentage of subjects reacting to peptides from each microorganism is variable but always <20% and the data from the THOR patients show significantly higher median values for all the microorganisms, in particular for Enterovirus B and C as well as Rhinovirus A (Fig. 7G). As for the comparison with the ELDERS, the median Log2fc change in the Ab titers reacting with the individual peptides from each microorganism is significantly lower in ADULTS than in THOR patients (Fig. 7 H).

Two peptides from HRSV are recognized by >50% of THOR patients and 22.2% of ADULTS with a relevant higher Ab titer (Log2fc >3) (Suppl. Table 5). Overall, the average percentage of responders in both groups is relatively low but significantly higher in thoracic patients (5.36% ADULTS; 14.35% THOR patients). Only 12 peptides are recognized by >20% of ADULTS (3 HHV-4, 2 HRSV, 2 Enterovirus B and C, 1 HHV-5, 1 HHV-1, 1 Staphylococcus aureus) and by a similar or higher percentage of THOR patients; on the contrary, 52 peptides are recognized by >20% of THOR patients and 28 of them (8 from HHV-5) are recognized by <10% of ADULTS. A significant number of peptides (38 peptides) are not recognized by any subject in the ADULTS group and 2 of them (pep_50960 from Adenovirus D; pep_115113 from Pegivirus A) are recognized by >20% of subjects in the THOR group. Similarly, 18 peptides are not recognized by any subject in the THOR group but the percentage of responders in the ADULTS group is very limited (<5%) (Suppl. Table 5).

### DSBP in HEALTHY (ELDERS and ADULTS) vs. SARC comparison

The comparisons between HEALTHY, ELDERS or ADULTS, and sarcoma patients (SARC) generated a DSBP pattern supporting the class prediction in both analyses, based on serological binding titers higher in the ELDERS vs. SARC and lower in the ADULTS vs. SARC comparisons (Fig. 8 A_1_ – A_3_, B_1_ – B_3_; Suppl. Table 6 and 7).

**Fig. 8.**
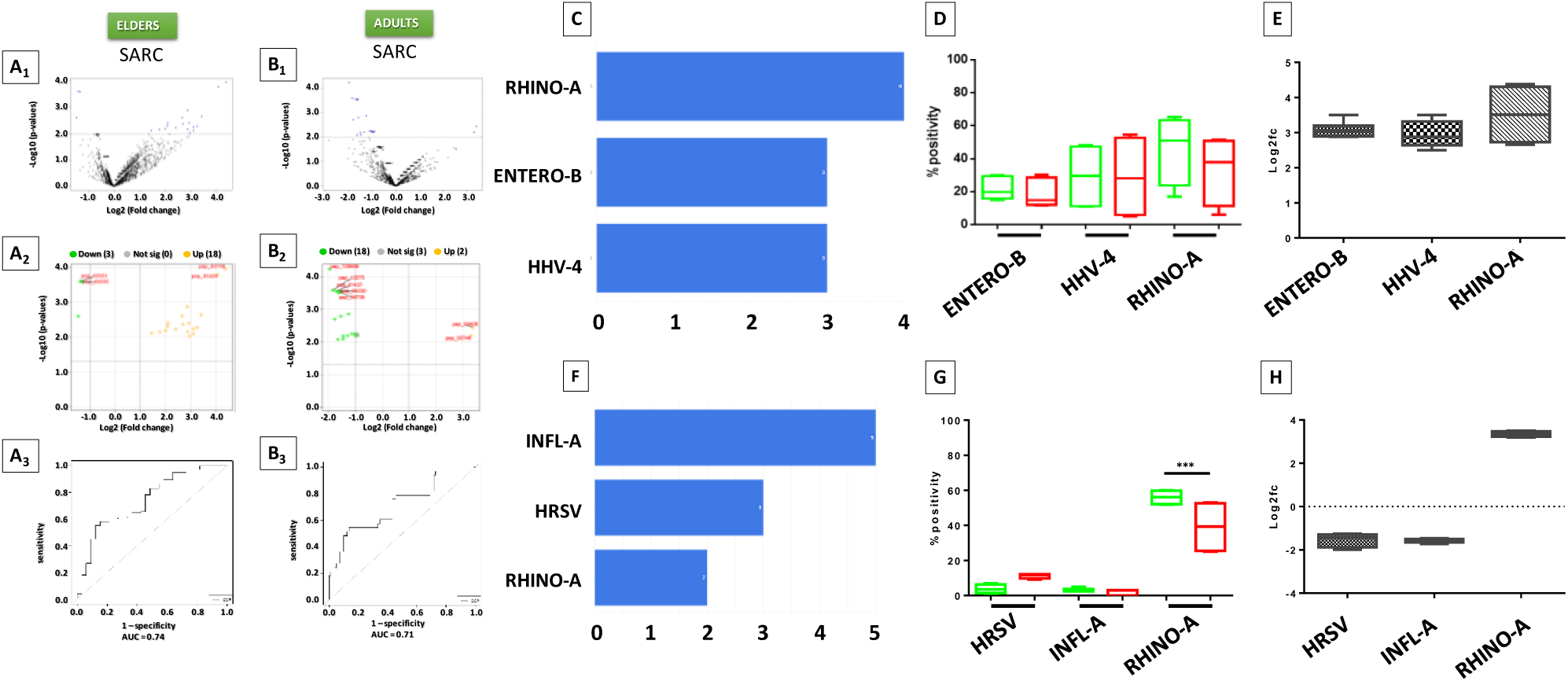
Differential serum binding pattern (DSBP) in HEALTHYvs.SARC comparison. Class comparison and prediction analysis based on antibody titers specific for the VirScan peptide array in ELDERS (A_1_ – A_3_) or ADULTS (B_1_ – B_3_) vs. SARC patients. Numerosity of peptides derived from the indicated microbes found in the DSBP (threshold of 2 peptides) (C and F). Percentage of responders in the two groups to all peptides from each microbial specie (D and G); differential binding titers between ELDERS (E) or ADULTS (H) vs. SARC patients.

The ELDERS vs. SARC DSBP is based on 18 peptides deriving from 13 microorganisms (all viruses) and the top three are Rhinovirus A (4 peptides), Enterovirus B and HHV-4 (3 peptides) (Fig. 8C). The median percentage of subjects reacting to peptides from each microorganism is significantly higher than the above-described comparisons but not statistically different in the two groups (Fig. 8D). Regardless the statistics, the Log2 fold change in the Ab titers reacting with the individual peptides from each microorganism, is highly relevant and higher in ELDERS than in SARC patients (Fig. 8E).

Two peptides from Rhinovirus A are recognized by >50% of ELDERS and by a similar percentage of SARC patients, likewise a single peptide from HHV-4 is recognized by >50% of SARC patients and by a similar percentage of ELDERS, in both cases this is supported by a relevant higher Ab titer (Log2fc ≥3) (Suppl. Table 6). A total of 8 peptides are recognized by >20% of ELDERS (3 Rhinovirus A, 2 HHV-4, 2 Enterovirus B, 1 HRSV and 1 HHV-1) and by a similar or higher percentage of SARC patients. Two peptides are not recognized by any subject in the SARC group and one (pep_111489 from Utive virus) is recognized by 13.2% of subjects in the ELDERS group (Suppl. Table 6).

The ADULTS vs. SARC DSBP is based on 20 peptides, of which 18 show a serological binding titer higher in SARC patients and 2 higher in ADULTS (Fig. 8B_1_ – B_3_). Notably, the latter are the same two peptides from the Rhinovirus A identified in the ELDERS vs. SARC comparison (Suppl. Table 7). Overall, the 20 peptides derive from 13 microorganisms (12 viruses) and the top three are Influenza A (5 peptides), HRSV (3 peptides) and Rhinovirus A (2 peptides) (Fig. 8F). The median percentage of ADULTS subjects and SARC patients reacting to peptides from each microorganism is variable but not statistically different and the Log2 fold change in the Ab titers shows that only for the Rhinovirus A it is higher in ADULTS than in SARC patients (Fig. 8G and H).

Overall, except for the high percentage of responders to the two Rhinoviruses peptides, the average percentage of responders in both groups is low (2.9% in ADULTS; 4.5% in SARC) and not statistically different. All peptides are recognized by ≤12% of subjects in both groups; a single peptide is not recognized by any subject in the ADULTS group (pep_31000 from Influenza B) and 6 peptides are not recognized by any subject in the SARC group. In all cases, the percentage of responders in the other group is very limited (≤7%) (Suppl. Table 7).

### Individual response to peptides involved the DSBP

The response to individual peptides from DSBP by each subject in the HEALTHY (ELDERS and/or ADULTS) and CANCER experimental cohort groups has been evaluated, showing two different outcomes: 1) peptides recognized by a different percentage of subjects in the two groups; and 2) peptides recognized by similar percentage of subjects in the two groups.

The DSBP in the comparisons between ELDERS and CANCER patients shows a higher percentage of responders in the ELDERS group to shared peptides from HRSV and HHV-5, especially when compared to sarcoma, breast, liver and melanoma patients. Even more evident is the significantly higher percentage of responders in the ELDERS group to individual peptides from microbes outside the most represented species (miscellaneous). Indeed, 11.9% of responses are observed in the ELDERS vs. 6% of all CANCER patients (3.5% without the THYR and THOR patients). Similar results are observed in the ELDERS vs. HCC comparison, with higher percentage of responders in the ELDERS group to peptides from HRSV and HHV-5 as well as from miscellaneous microbes, namely 13.3% of responses observed in the ELDERS vs. 2.5% of HCC patients.

Otherwise, the percentage of responders to peptides from other persistent herpesviruses is not different in subjects from the experimental groups with an antibody titer statistically higher in ELDERS (p-value <0.01) (Fig. 9A) (Suppl. Fig. 3).

**Fig. 9.**
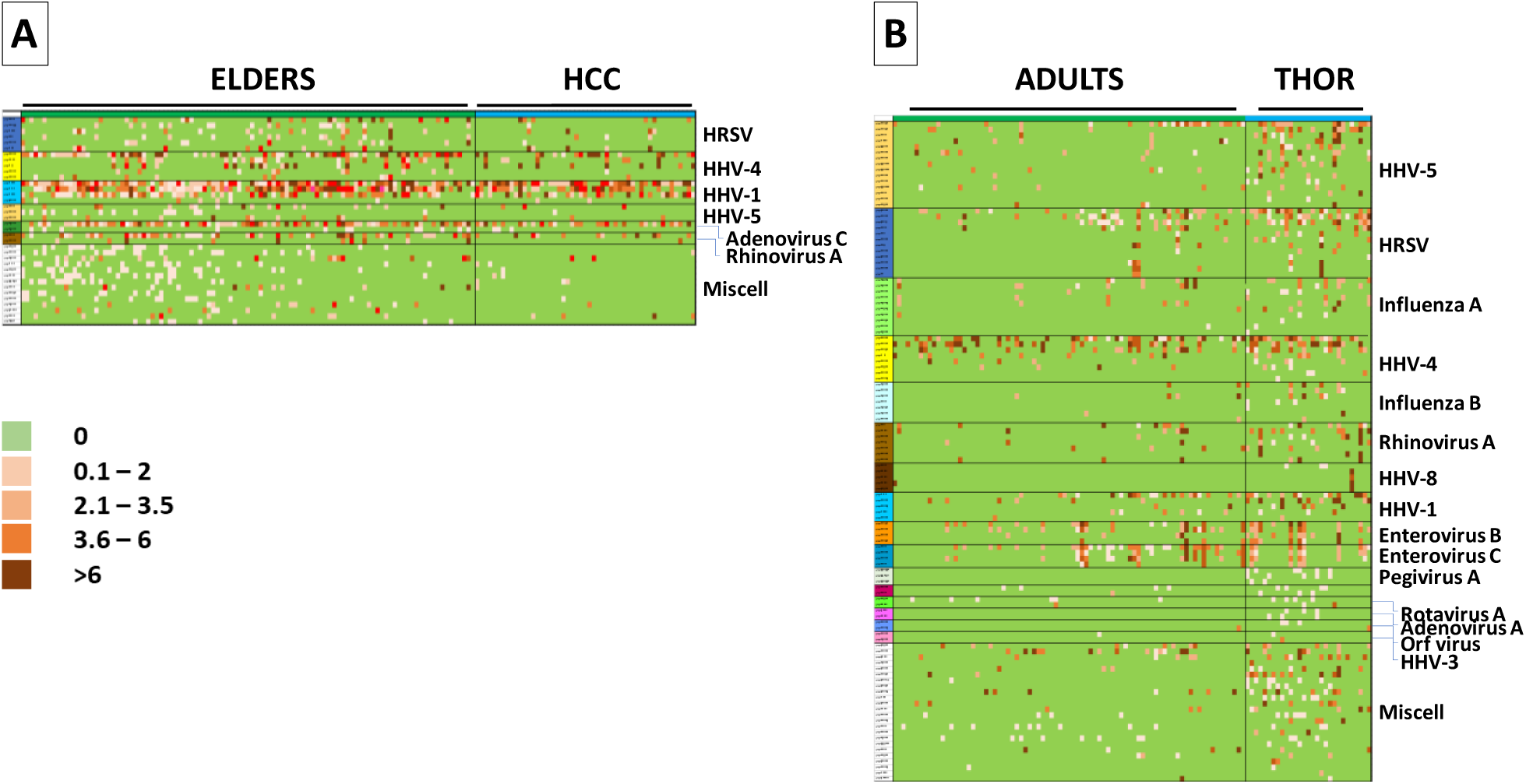
Map of responders to peptides in HEALTHY vs. CANCER comparison. Responders to each peptide from the indicated microbe species are shown as heatmaps, ELDERSvs.HCC (A) ADULTSvs.THOR (B). Miscell = peptides identified from single microbes.

The DSBP in the comparison between ADULTS and all CANCER patients shows a higher percentage of responders in the CANCER group to shared peptides from HRSV, while the percentage of responders to shared peptides from Influenza A, Enterovirus B, HHV-5 and miscellaneous is comparable between groups.

On the contrary, the one-to-one comparison with THOR and THYR patients shows a more articulated picture. Overall, the higher percentage of responders in THOR and THYR patients is observed to peptides from several microbes, including persistent herpesviruses and acute-infecting (HRSV, Influenza A and B, Rhinovirus A) viruses. Interestingly, THYR and THOR patients respond to the same peptides with a statistically higher antibody titer than ADULTS and the higher percentage of responders is observed to peptides from HRSV. Similar to the data observed in the ELDERS vs. CANCER comparison, a higher percentage of responders is found to peptides from miscellaneous microbes. However, in this case, CANCER patients are more responders than ADULTS. Indeed, 15.1% of responses are observed in the THOR patients vs. 3.9% of ADULTS; 14.2% of responses are observed in the THYR patients vs. 1.03% of ADULTS.

The percentage of responders to peptides from other microbes is not different in subjects from the experimental groups, although the antibody titer is statistically higher in CANCER groups (p-value <0.01). Of note, besides chronic herpesviruses, peptides from several acute viruses (including, Enteroviruses B and C, Influenza A and B, Rhinovirus A) play a key role in the DSBP predicting ADULTS and THOR & THYR patients (Fig. 9B) (Suppl. Fig. 4).

### Pool of peptides classifying HEALTHY and CANCER subjects

To identify a specific peptide panel for the diagnostic discrimination between healthy individuals and cancer patients, the peptides were selected if the responses were significantly divergent the two groups (namely ≤5% in a group and ≥20% in the other group).

The results showed two distinctive patterns when considering the responses in ≤5% of cancer patients or in healthy individuals.

The peptides with responses in ≤5% of cancer patients (min 0% – max 5%, median value 1.15%) and ≥20% of healthy individuals (min 20% – max 62.26%, median value 25.47%) predominantly derive from HHV-5. Looking into the ELDERS and ADULTS groups independently, there is an astounding difference. Indeed, the responses in ELDERS are to 53 peptides predominantly from several persistent herpesviruses and, at a lower percentage, from HRSV, Influenza A and B as well as Orf virus. Of note, the response to two peptides from HHV-5 is 0.0% in melanoma patients and 58.5-62.2% in ELDERS (Suppl. Fig. 5). On the contrary, the responses in ADULTS are to 7 peptides predominantly from Poliovirus (4 peptides) and HHV-5 (2 peptides); the latter two peptides are shared with the ELDERS (Suppl. Fig. 5) (Fig. 10 A – E).

**Fig. 10.**
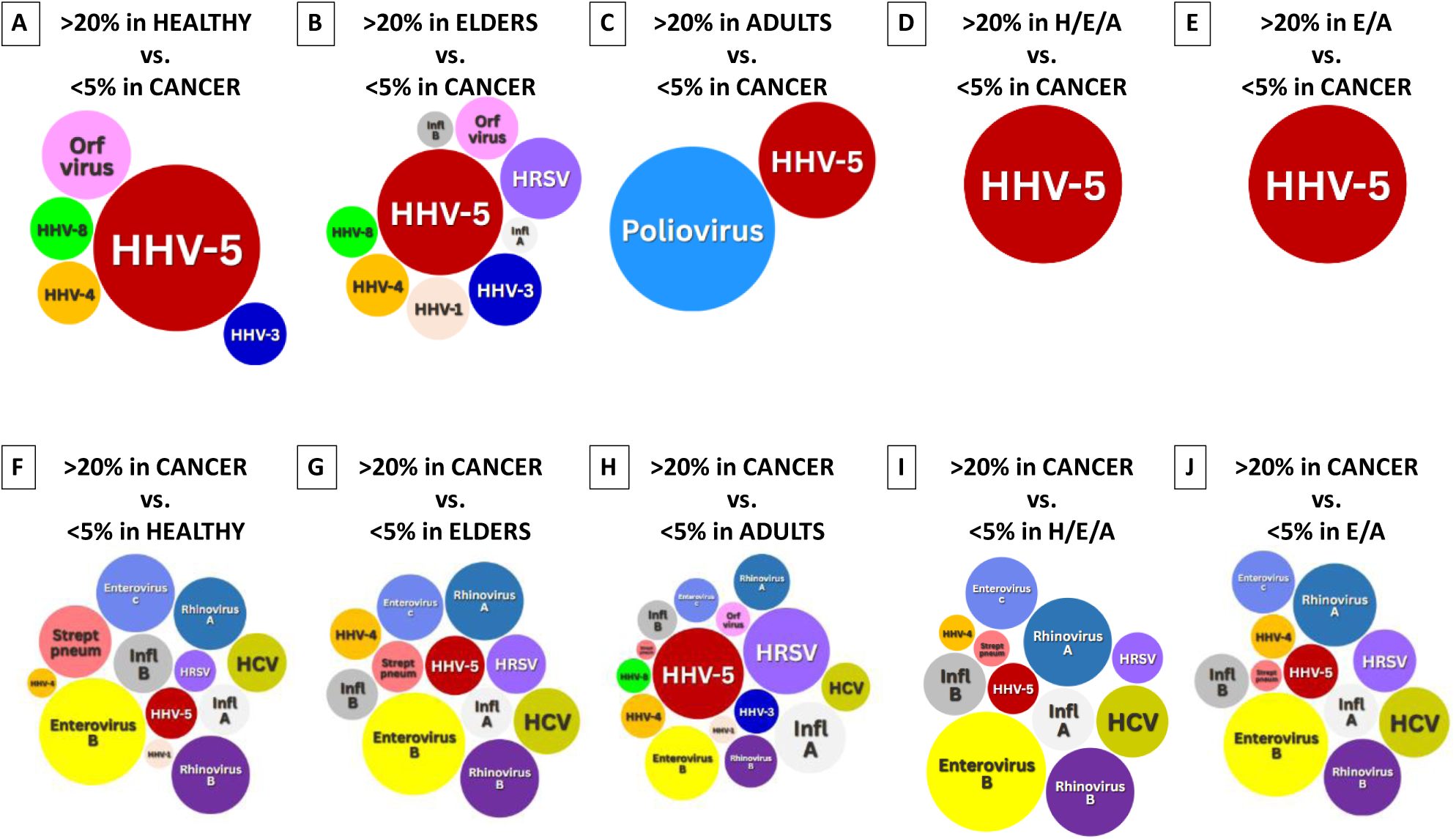
Peptides classifying HEALTHY and CANCER subjects. Numerosity of microbes from which are derived the peptides with >20% of responders in HEALTHY (or ELDERS or ADULTS) and <5% in CANCER subjects (A – E). Numerosity of microbes from which are derived the peptides with <5% of responders in HEALTHY (or ELDERS or ADULTS) and >20% in CANCER subjects (F – J).

The peptides with responses in ≤5% of healthy individuals (min 0% – max 5%, median value 2.47%) and ≥20% of cancer patients (min 20% – max 62.15%, median value 23.53%) mostly derive from acute infecting agents, prevalently Enterovirus B and C as well as Rhinoviruses A and B. HHV-5 is the only persistent herpesvirus present in this analysis, but the number of peptides is very limited. In this case, the independent evaluation of ELDERS and ADULTS groups showed a remarkable overlapping with a limited overrepresentation of peptides from HHV-5 and HRSV in the comparison with ADULTS individuals (Suppl. Fig. 6) (Fig. 10 F – J). In particular, the combined comparison to both ELDERS and ADULTS showed 71 shared peptides with a low response in HEALTHY individuals (median 2.45%) and a high in CANCER patients (median 23.08%). Of note, the response to one peptide from Enterovirus B is 3.5% in both HEALTHY groups and 31% (thoracic patients) and 54.5% (melanoma patients) (Suppl. Fig. 6).

## Discussion

Sera from two experimental groups, HEALTHY and CANCER subjects, have been collected for assessing the antibody titers specific to 82,134 peptides covering the entire spectrum of human virome and bacteriome. The age in the cancer cohort shows a unimodal distribution with a median at 64.07 years, while the healthy cohort shows a bimodal distribution with a median at 50.3 and at 88.1 years, respectively. This identifies two sub-cohorts of volunteers named ADULTS (<70 years of age) and ELDERS (>80 years of age). Overall, sera from both HEALTY and CANCER cohorts show a reactivity to 39.3% of the peptides, while none of the subjects in both groups show reactivity against 70,300 peptides (60.7%).

The top peptide is derived from Streptococcus pneumoniae, with >97% of subjects from both groups showing antibody reactivity. The peptides with >50% reacting healthy subjects (nr. 54) derive from 8 pathogens and the top three are HHV-1 (15 peptides), HRSV (9 peptides), Rhinovirus-A (8 peptides). Among these 54 peptides, a significant lower percentage of reactivity (<35%) is observed in the CANCER group for 2 peptides from HHV-4 and 1 from HHV-1. The class comparison performed with serological data obtained in all cancer patients vs. the ELDERS or ADULTS subjects, separately, show a differential serum binding pattern (DSBP) supporting a class prediction, as confirmed by the receiver operating characteristic (ROC) curve analysis. Interestingly, while the DSBP of the ELDERS vs. CANCER comparison is characterized by higher titers in the healthy subjects, the DSBP of the ADULTS vs. CANCER comparison is characterized by lower titers in the healthy subjects. Regardless the symmetrical pattern of Ab titers, it is notable that the highest number of peptides identified in the two DSBP are from HRSV. Besides such a predominance, the DSBP supporting the ELDERS class prediction is mostly based on higher Ab titers to herpesviruses in healthy subjects, while the one supporting the ADULTS class prediction is mostly based on higher Ab titers to acute viruses in cancer patients (e.g. influenza A and Enterovirus B).

The same analysis performed comparing the healthy subjects to each subgroup of cancer patients generated quite different outcomes, with DSBP supporting a class prediction only in selected comparisons. In particular, predictive patterns are found when 1) ELDERS are compared to HCC patients; 2) ADULTS are compared to THYROID ca patients; 3) ELDERS or ADULTS are compared to SARCOMA patients; and 4) ELDERS or ADULTS are compared to THORACIC ca patients. Interestingly, all these class predictions are based on the same pattern characterized by higher titers in ELDERS and lower titers in ADULTS compared to CANCER patients.

All the DSBP supporting the individual class predictions are mainly or mostly based on differential GMT of Ab responses to several peptides derived from persistent herpesviruses (namely HHV-4, HHV-5 and HHV-1). In addition to these, a consistent pattern is the differential responses to several peptides from acute-infecting viruses, prevalently Influenza A, Enterovirus B and C, Rhinovirus A. Strikingly, except for the comparison with the SARC patients, the highest number of peptides identified in the DSBP are derived from the HRSV. The percentage of responders to each peptide characterizing the individual DSBP is highly variable in the comparisons and three conditions of percentage of responders between the HEALTHY and CANCER groups may be classified: 1) the percentage is not significantly different; 2) the percentage is significantly different; and 3) the percentage is zero in individuals (no response in any individual) of one of the two groups. Overall, the Log2fc in the GMT is always >1, reaching also a value >5, and the statistical significance is supported by a p-value <0.01.

In the ELDERS vs. HCC comparison there is a significant higher percentage of responders in the healthy subjects to peptides from HHV-5, HRSV and Enterovirus B. Collectively, several individual peptides are differentially recognized, but only two (from Rhinovirus A and HHV-4) are above a threshold of 50% of responsive subjects (ELDERS>HCC). Few peptides are not recognized by HCC patients but the percentage of responders in the ELDERS group is <20%.

In the ADULTS vs. THYR comparison there is a significant lower percentage of responders in the healthy subjects to peptides from Enterovirus B and C as well as Influenza A. In particular, several peptides from Influenza A are recognized by >20% of THYR patients a <5% (or even 0%) of ADULTS subjects. Strikingly, a single peptide from the Staphylococcus Aureus is recognized by about 70% of THYR patients and <30% of ADULTS individuals as well as a single peptide from Adenovirus C is recognized by about 46% of THYR patients and 1.2 % of ADULTS individuals.

In the HEALTHY (ELDERS and ADULTS) vs. THOR comparison there is a significant lower percentage of responders in the healthy subjects to peptides from Enterovirus B, Rhinovirus A and B (both ELDERS and ADULTS), as well as Adenovirus A, HRSV and HHV-5 (ADULTS). Interestingly, few peptides are in common to both comparisons from HHV-4, Adenovirus A, HHV-6A, Enterovirus B and C. In the ADULTS vs. THOR comparison, several peptides from different organisms (e.g. Influenza A, Enterovirus B and C, HHV-4, HHV-1, HHV-5, HRSV, Rhinovirus A) are recognized by >20% of THOR patients a <10% (or even 0%) of ADULTS subjects. Strikingly, two peptides from HRSV are recognized by about >50% of THOR patients and 22.2% of ADULTS individuals.

The heatmaps of the different qualitative and quantitative responses in each subject to individual peptides from the different microorganisms confirmed the DSBP, identifying a selected number of peptides differentially recognized by a relevant percentage of subjects in the HEALTHY (ELDERS or ADULTS) subjects or CANCER patients. In particular, considering the entire set of responses in the different comparisons, a handful of epitopes strongly characterize the two groups of enrolled subjects and allow a conclusive classification.

Filtering the data on significantly divergent responses to peptides in the two groups (namely ≤5% in a group and ≥20% in the other group), the results are strikingly different. Indeed, the peptides with ≤5% of responses in CANCER patients and ≥20% in HEALTHY individuals are mostly from persistent herpesviruses (with a predominance of HHV-5), in ELDERS, and Poliovirus and HHV-5, in ADULTS. On the contrary, the source of peptides with ≤5% of responses in HEALTHY individuals and ≥20% in CANCER patients is completely different, with an overwhelming representation of acute infecting microbes.

Overall, the higher percentages in ELDERS of responses to peptides derived from herpesviruses, prevalently HHV-5, is largely known due to lifelong repeated viral reactivation and boosting of the immune system (27). In parallel, the response to novel acute infecting pathogens and antigens has been reported to be reduced as result of the immunosenescence with the progressive loss of naïve B cells (28). This should result in an impaired adaptability and robustness of immune defenses in older individuals and concomitant increased incidence of cancer (29–31). Nonetheless, in the present study, a significant number of responses to peptides derived from acute infecting viruses was significantly higher in healthy ELDERS vs. CANCER patients, namely HRSV, Influenza A and B, Chikungunya. In parallel, ADULTS are classified based on higher responses to two HHV-5 peptides (shared with ELDERS) and, uniquely, to four peptides derived from poliovirus type 1 (>25% of responders) which are recognized by <4% of cancer patients (namely HCC and HNSCC). Such results are mirrored by the finding of significantly higher responses in CANCER patients vs. ELDERS or ADULTS to several shared peptides derived by a broad spectrum of acute infecting pathogens. Strikingly, these are prevalently Enteroviruses B and C, Rhinoviruses A and B, Influenza viruses and HRSV. Interestingly, the spectrum of pathogens is almost identical in the comparisons with ELDERS or ADULTS, normalizing the bias due to the different ages of the two subgroups of healthy individuals. Such results are in agreement to previous reports showing the key role of antibody responses to such acute infecting pathogens in a comparative analysis focused on healthy individuals and HCC patients (32).

The present study represents a proof of concept to be extended on a larger population covering broader geographical regions. In the present design, it showed novel supportive data about a significant correlation between differential serum binding patterns (DSBP) in healthy individuals and cancer patients, providing significant experimental evidences of responses to specific microbe-derived peptides valuable for a clinical classification. Notably, such evidences disclosed binding patterns shared by ADULTS (<70 years of age) and ELDERS (>80 years of age) when compared to CANCER patients, suggesting the possibility of selecting a pool of peptides as biomarkers for prediction and/or diagnostic tool.

## Limitations of the study

Our finding that responses to acute-infecting viruses are significantly different in healthy individuals and cancer patients could be influenced by several confounding factors, including transmissibility (33) and socioeconomic conditions (34). There are also several important limitations related to the cohort design. Indeed, the study has been conducted in a single Italian Region (Campania) and volunteers have been enrolled in a single large city hospital center (ADULTS and CANCER patients) or in villages from a single inland area (ELDERS). The latter were purposely selected as example of “real” tumor-free individuals with an age >80 yo. Regardless such a controlled bias, cancer patients showed higher responses to the same acute-infecting agents in the comparison with both ELDERS and ADULTS.

Finally, the technology does not allow to distinguish between more recent infection or long-term immunity, due to virus-dependent differential persistence of serum antibodies.

In future research efforts, the present data will be examined in prospective studies on a broader cohort population from a larger geographical area in order to mitigate the evidenced limitations of the present study and support the associations herein described.

## Legenda

HHV-1-8: Human Herpesvirus 1-8
HRSV: Human Respiratory Syncytial Virus
Rhino: Human Rhinovirus
Entero: Human Enterovirus
HCV: Hepatitis C virus
HBV: Hepatitis B virus
HEV: Hepatitis E virus
INFL: Influenza virus
ADENO: Human adenovirus
Entam Hist: Entamoeba histolytica
Mollusc Cont: Molluscum contagiosum
HCC: hepatocellular carcinoma
BREA: breast cancer
SARC: sarcoma
THOR: Lung cancer
THYR: thyroid cancer
MEL: melanoma
HNSCC: head&neck sqamous cell carcinoma

## Acknowledgements

We sincerily thank all the family doctors who were actively involved in the recruitment of ELDERS at the villages of the the “Comunità Montanta del Fortore”, Benevento – IT. Furthermore, we are indebited to all the healthy subjects and cancer patients, in particular the ELDERS, who accepted to participate.

## Declarations

### 1. Ethics approval and consent to participate

The study has been approved by the Institutional Ethical Committee of the Istituto Nazionale Tumori - IRCCS - “Fond G. Pascale” – NAPOLI IT (Protocol n. 57/22 OSS “HepAnt – Tumor antigen discovery for innovative cancer immunotherapies in HCC: from benchside to bedside” e subsequent amendments)

### 2. Consent for publication

The corresponding author has received consent for publication.

### 3. Availability of data and material

Data and material will be deposited and publicly available.

### 6. Authors’ contributions

BC performed all the bioinformatics analyses; BC and SM processed all the serological samples; AM and CR contributed to the generation and maintenance of the database; LW, CM, YZ and XWW performed and validated the VirScan analysis; AB, FI, RD’A, MGC, CAF, AM, AC, MDL, VV, PA, FP, OC, NM, EP, EM and FI enrolled the subjects and provided the blood samples; MT and LB designed and coordinated the study; BC, MT and LB drafted the manuscript. All Authors approved the manuscript.

