## Supplementary material for "Differential serum binding patterns predicting healthy subjects and cancer patients": Suppl. figures

### Suppl. Fig. 1

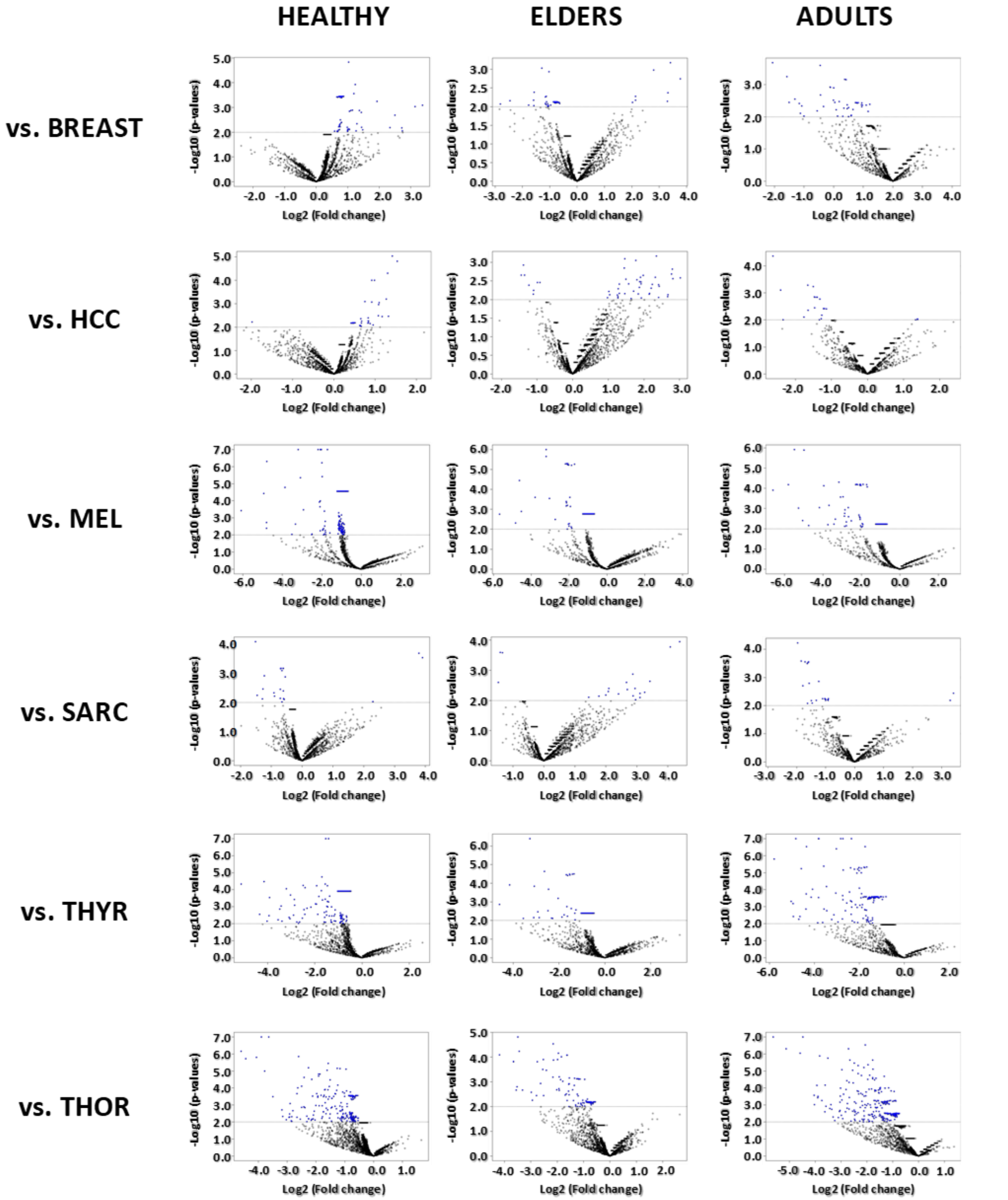


Suppl. Fig. 1 Class comparison analysis based on antibody titers specific for the VirScan peptide array. HEALTHY, or ELDERS or ADULTS subjects vs. CANCER patients with different diagnosis.

### Suppl. Fig. 2

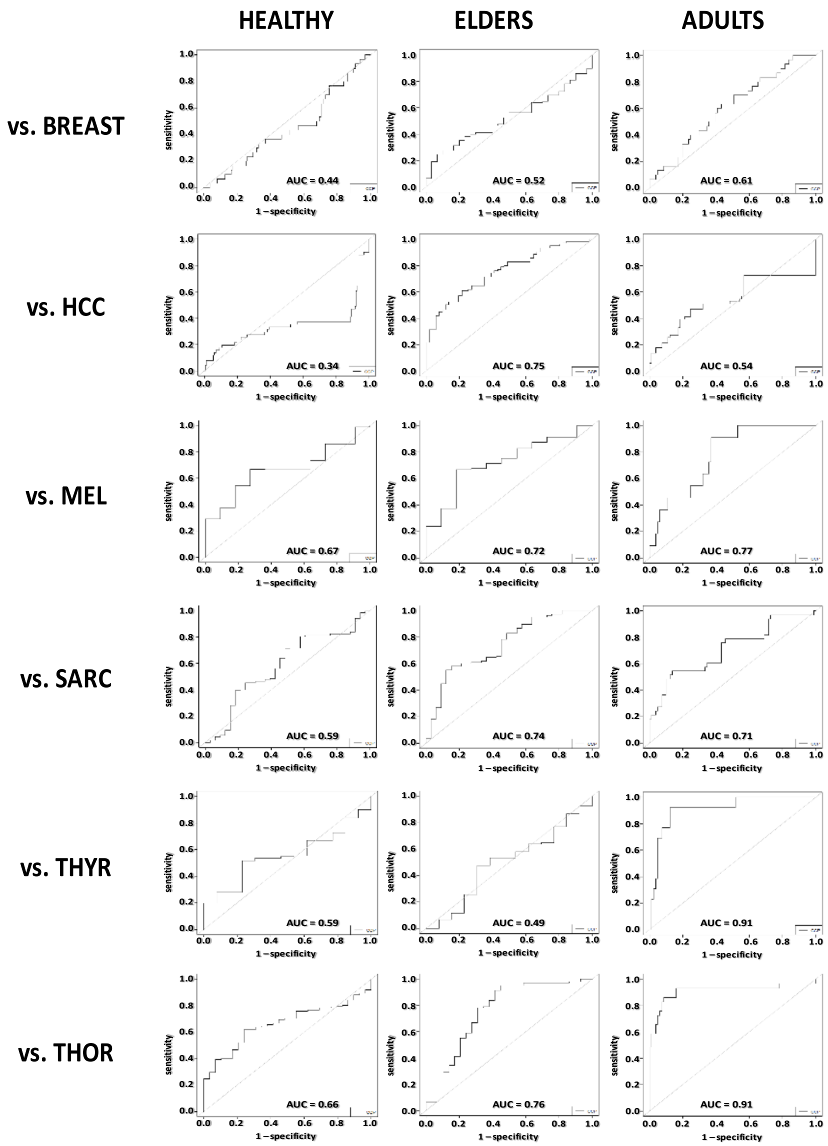


Suppl. Fig. 2 Class prediction analysis based on antibody titers specific for the VirScan peptide array. HEALTHY, or ELDERS or ADULTS subjects vs. CANCER patients with different diagnosis. The ROC curves derived from class prediction analysis for each of the comparisons are shown. AUC = area under the curve.

### Suppl. Fig. 3

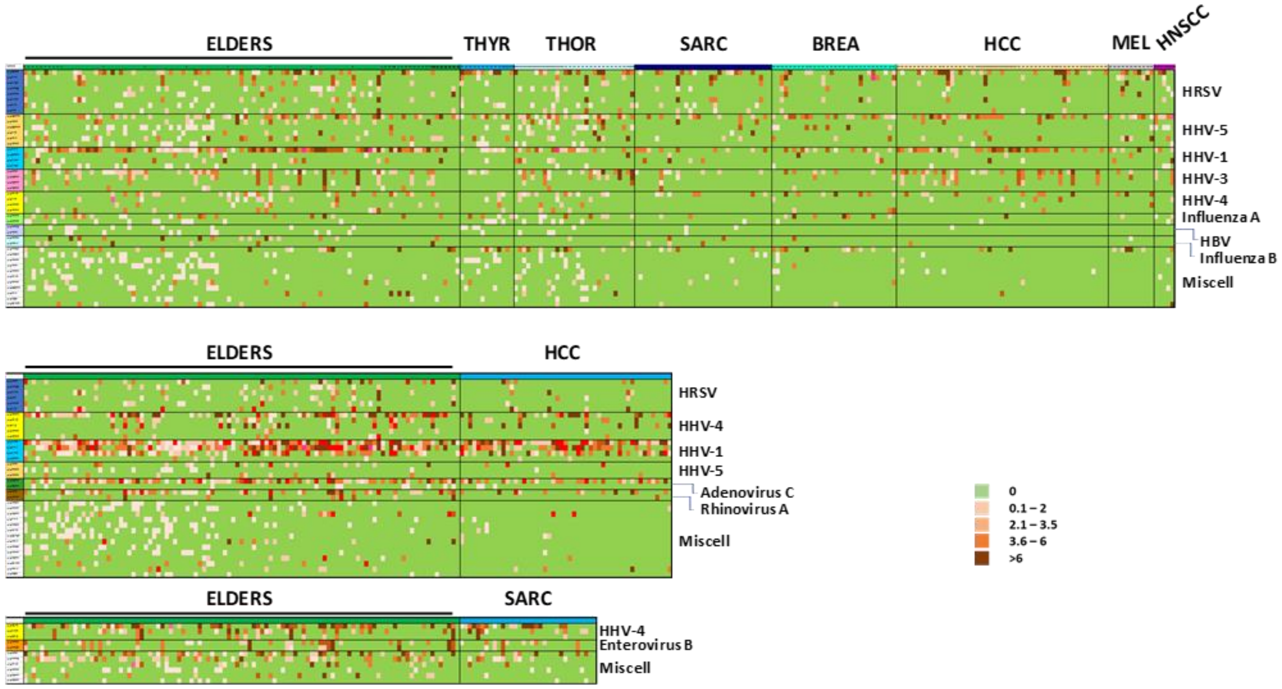


Suppl. Fig. 3 Map of responders to peptides in ELDERSvs.CANCER comparison. Responders to each peptide from the indicated microbe species are shown as heatmaps. Miscell = peptides identified from single microbes.

#### Suppl. Fig. 4

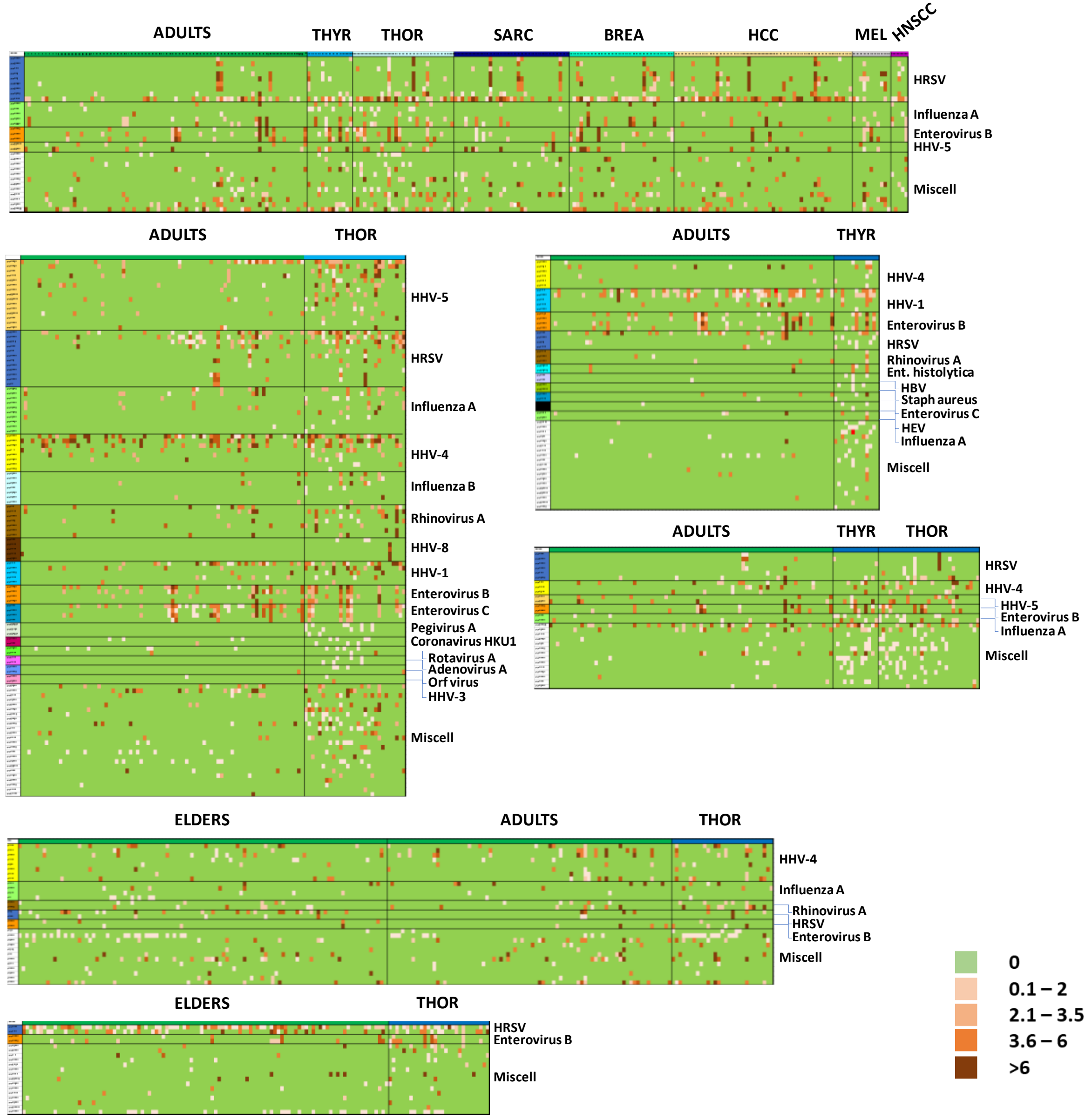


Suppl. Fig. 4 Map of responders to peptides in ADULTSvs.CANCER comparison. Responders to each peptide from the indicated microbe species are shown as heatmaps.
Miscell = peptides identified from single microbes

**Suppl. Fig. 5**

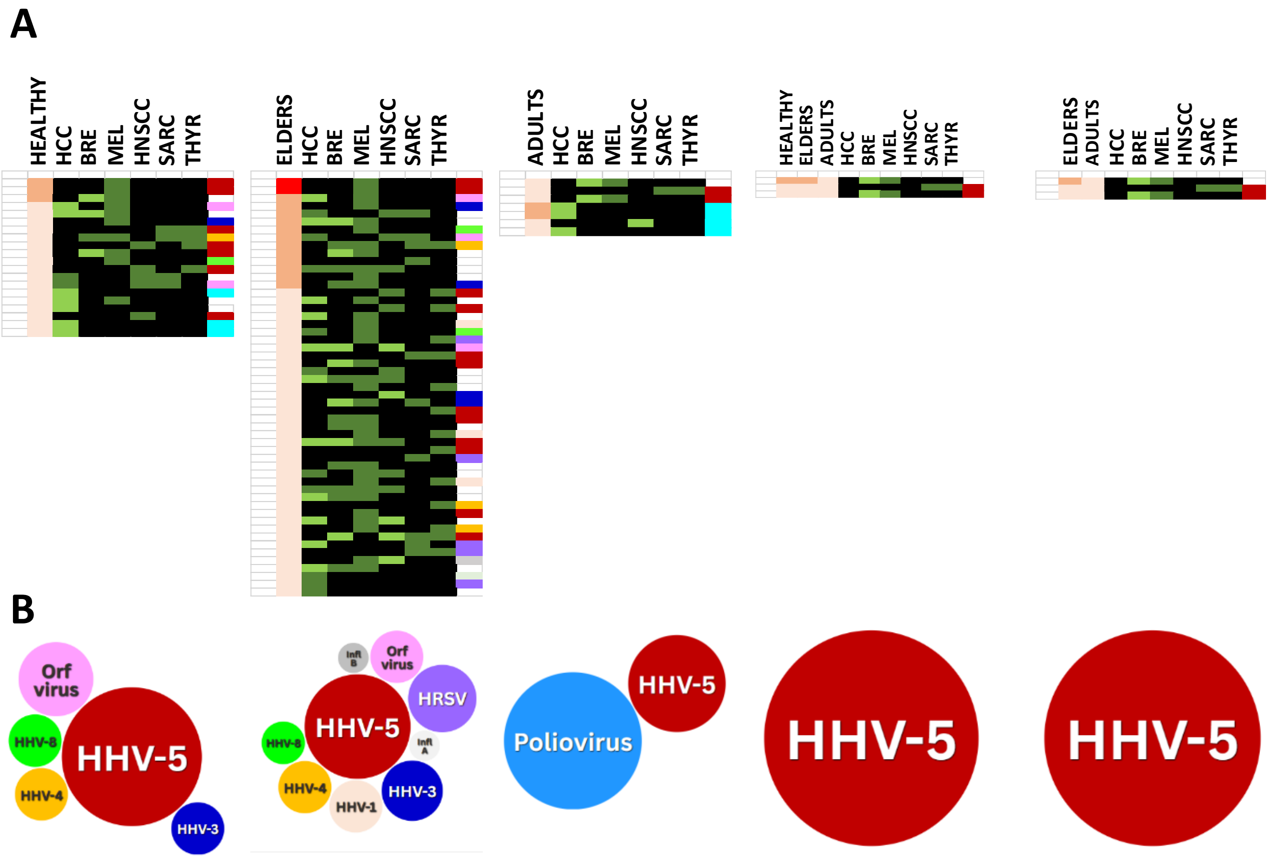


Suppl. Fig. 5 Peptides classifying HEALTHY and CANCER subjects. (A) Individual peptides with higher number of responders in HEALTHY (or ELDERS or ADULTS) individuals (>20%) than in CANCER patients (<5%). (B) Numerosity of microbes from which are derived the peptides.

### Suppl. Fig. 6

A

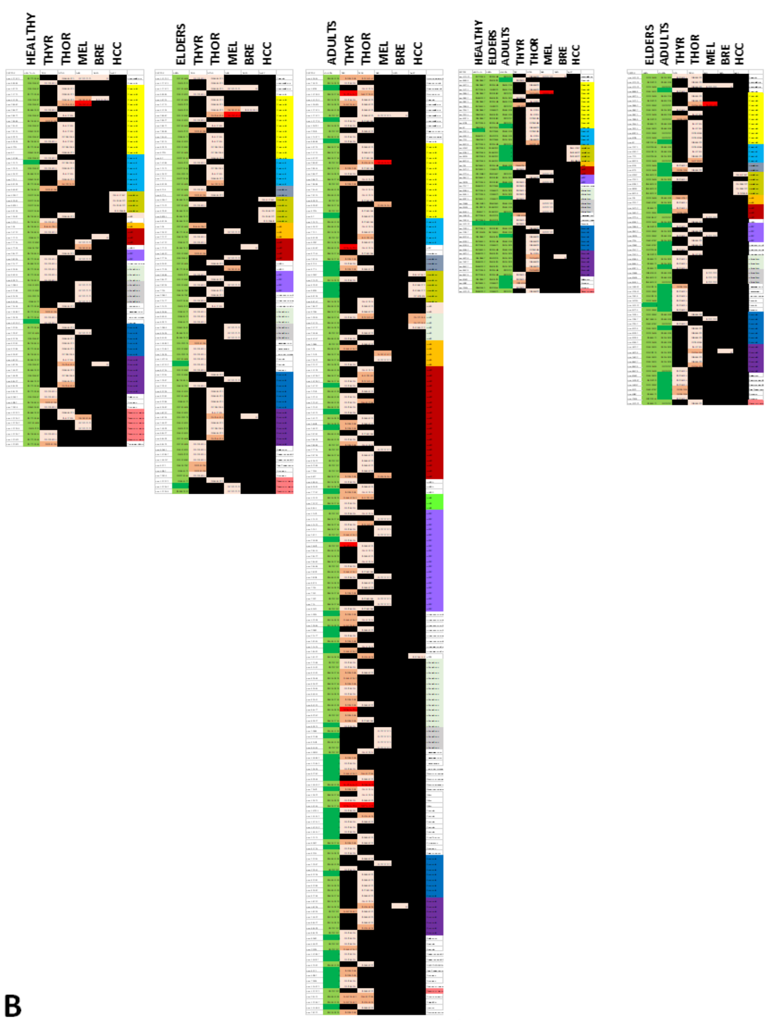


B

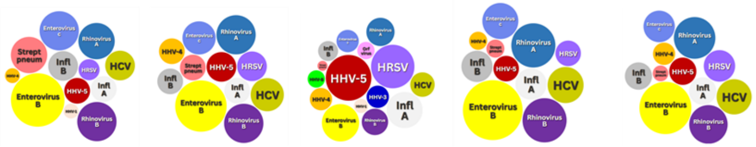


Suppl. Fig. 6 Peptides classifying HEALTHY and CANCER subjects. (A) Individual peptides with lower number of responders in HEALTHY (or ELDERS or ADULTS) individuals (<5%) than in CANCER patients (>20%). (B) Numerosity of microbes from which are derived the peptides.
