## Supplementary material for "Differential serum binding patterns predicting healthy subjects and cancer patients": Suppl. Tables

**Suppl. Table 1.** Peptides identified in the differential serum binding pattern (DSBP) in the comparison between ELDERS or ADULTS and all CANCERS with percentage of responders.

| ID | ELDERS | THOR | SARC | MEL | BRE | HEP | THYR |
|---|---|---|---|---|---|---|---|
| 115299 | 43,40% | 68,97% | 30,30% | 63,64% | 56,67% | 47,06% | 53,85% |
| 30608 | 77,36% | 58,62% | 30,30% | 36,36% | 36,67% | 60,78% | 53,85% |
| 13123 | 37,74% | 48,28% | 36,36% | 54,55% | 30,00% | 41,18% | 76,92% |
| 85617 | 16,04% | 31,03% | 12,12% | 45,45% | 33,33% | 5,88% | 38,46% |
| 25890 | 45,28% | 51,72% | 24,24% | 18,18% | 23,33% | 43,14% | 30,77% |
| 37187 | 15,09% | 13,79% | 6,06% | 0,00% | 3,33% | 1,96% | 7,69% |
| 95657 | 18,87% | 27,59% | 6,06% | 9,09% | 23,33% | 1,96% | 30,77% |
| 56034 | 21,70% | 34,48% | 21,21% | 27,27% | 23,33% | 15,69% | 15,38% |
| 51732 | 17,92% | 31,03% | 6,06% | 9,09% | 6,67% | 21,57% | 0,00% |
| 56780 | 16,04% | 27,59% | 9,09% | 0,00% | 0,00% | 5,88% | 0,00% |
| 115293 | 19,81% | 37,93% | 3,03% | 9,09% | 23,33% | 0,00% | 23,08% |
| 70430 | 11,32% | 13,79% | 12,12% | 36,36% | 23,33% | 9,80% | 15,38% |
| 70418 | 14,15% | 13,79% | 9,09% | 9,09% | 13,33% | 11,76% | 15,38% |
| 20950 | 12,26% | 10,34% | 0,00% | 0,00% | 3,33% | 9,80% | 15,38% |
| 56411 | 16,98% | 24,14% | 9,09% | 9,09% | 10,00% | 1,96% | 0,00% |
| 55858 | 9,43% | 10,34% | 0,00% | 9,09% | 0,00% | 1,96% | 7,69% |
| 56495 | 12,26% | 6,90% | 0,00% | 18,18% | 3,33% | 1,96% | 0,00% |
| 15236 | 6,60% | 3,45% | 3,03% | 9,09% | 6,67% | 3,92% | 0,00% |
| 51737 | 14,15% | 10,34% | 3,03% | 0,00% | 0,00% | 21,57% | 0,00% |
| 70425 | 14,15% | 10,34% | 12,12% | 0,00% | 13,33% | 7,84% | 15,38% |
| 59541 | 9,43% | 3,45% | 6,06% | 0,00% | 0,00% | 1,96% | 30,77% |
| 22694 | 13,21% | 10,34% | 3,03% | 0,00% | 3,33% | 1,96% | 46,15% |
| 28530 | 13,21% | 10,34% | 0,00% | 0,00% | 3,33% | 0,00% | 7,69% |
| 83385 | 14,15% | 3,45% | 6,06% | 9,09% | 3,33% | 0,00% | 0,00% |
| 7043 | 10,38% | 17,24% | 18,18% | 0,00% | 0,00% | 0,00% | 30,77% |
| 27662 | 9,43% | 6,90% | 0,00% | 0,00% | 6,67% | 7,84% | 7,69% |
| 3535 | 10,38% | 10,34% | 3,03% | 0,00% | 0,00% | 0,00% | 7,69% |
| 2940 | 13,21% | 17,24% | 12,12% | 0,00% | 3,33% | 5,88% | 0,00% |
| 8083 | 11,32% | 13,79% | 6,06% | 0,00% | 0,00% | 1,96% | 0,00% |
| 95097 | 13,21% | 20,69% | 6,06% | 0,00% | 0,00% | 0,00% | 7,69% |
| 9607 | 21,70% | 17,24% | 9,09% | 36,36% | 10,00% | 7,84% | 30,77% |
| 80496 | 13,21% | 17,24% | 12,12% | 9,09% | 6,67% | 3,92% | 7,69% |
| 61544 | 10,38% | 3,45% | 3,03% | 0,00% | 3,33% | 1,96% | 0,00% |
| 34232 | 13,21% | 13,79% | 0,00% | 0,00% | 0,00% | 1,96% | 23,08% |
| 83955 | 11,32% | 17,24% | 3,03% | 0,00% | 3,33% | 1,96% | 30,77% |
| 48897 | 11,32% | 10,34% | 6,06% | 0,00% | 3,33% | 1,96% | 7,69% |
| 111489 | 13,21% | 20,69% | 0,00% | 0,00% | 3,33% | 0,00% | 15,38% |
| 60316 | 11,32% | 6,90% | 0,00% | 0,00% | 3,33% | 1,96% | 0,00% |
| 52988 | 9,43% | 10,34% | 0,00% | 0,00% | 10,00% | 1,96% | 7,69% |
| 65050 | 6,60% | 10,34% | 0,00% | 0,00% | 0,00% | 1,96% | 0,00% |
| 9115 | 8,49% | 13,79% | 0,00% | 0,00% | 0,00% | 0,00% | 7,69% |
| 120234 | 9,43% | 3,45% | 3,03% | 0,00% | 3,33% | 0,00% | 0,00% |
| 83407 | 6,60% | 3,45% | 9,09% | 0,00% | 0,00% | 1,96% | 0,00% |
| 8005 | 2,83% | 10,34% | 12,12% | 0,00% | 0,00% | 0,00% | 0,00% |

| ID | ADULTS | THOR | SARC | MEL | BRE | HEP | THYR |
|---|---|---|---|---|---|---|---|
| 82438 | 4,9% | 17,2% | 3,0% | 0,0% | 0,0% | 0,0% | 7,7% |
| 125606 | 1,2% | 0,0% | 6,1% | 0,0% | 10,0% | 9,8% | 7,7% |
| 22259 | 1,2% | 6,9% | 12,1% | 0,0% | 3,3% | 7,8% | 7,7% |
| 58427 | 2,5% | 17,2% | 9,1% | 0,0% | 0,0% | 2,0% | 7,7% |
| 16322 | 3,7% | 10,3% | 9,1% | 9,1% | 10,0% | 9,8% | 15,4% |
| 95377 | 7,4% | 10,3% | 0,0% | 9,1% | 16,7% | 2,0% | 23,1% |
| 25542 | 2,5% | 10,3% | 12,1% | 0,0% | 0,0% | 3,9% | 0,0% |
| 7028 | 1,2% | 10,3% | 9,1% | 0,0% | 3,3% | 0,0% | 0,0% |
| 731 | 4,9% | 10,3% | 9,1% | 27,3% | 16,7% | 7,8% | 15,4% |
| 23621 | 8,6% | 27,6% | 6,1% | 0,0% | 10,0% | 3,9% | 0,0% |
| 2510 | 4,9% | 13,8% | 12,1% | 27,3% | 23,3% | 15,7% | 23,1% |
| 92942 | 4,9% | 10,3% | 3,0% | 0,0% | 10,0% | 3,9% | 30,8% |
| 13113 | 4,9% | 13,8% | 12,1% | 27,3% | 13,3% | 13,7% | 7,7% |
| 12275 | 3,7% | 10,3% | 12,1% | 18,2% | 10,0% | 13,7% | 7,7% |
| 115288 | 7,4% | 10,3% | 6,1% | 9,1% | 6,7% | 2,0% | 0,0% |
| 11867 | 6,2% | 10,3% | 12,1% | 27,3% | 10,0% | 13,7% | 15,4% |
| 94230 | 2,5% | 20,7% | 3,0% | 0,0% | 0,0% | 0,0% | 30,8% |
| 94735 | 2,5% | 10,3% | 0,0% | 9,1% | 3,3% | 3,9% | 0,0% |
| 85871 | 11,1% | 27,6% | 3,0% | 27,3% | 13,3% | 2,0% | 30,8% |
| 72527 | 4,9% | 17,2% | 9,1% | 0,0% | 3,3% | 3,9% | 23,1% |
| 19082 | 7,4% | 31,0% | 21,2% | 18,2% | 26,7% | 13,7% | 7,7% |
| 115280 | 8,6% | 20,7% | 18,2% | 18,2% | 23,3% | 11,8% | 30,8% |
| 81422 | 11,1% | 17,2% | 9,1% | 18,2% | 20,0% | 5,9% | 30,8% |
| 81114 | 11,1% | 27,6% | 15,2% | 18,2% | 20,0% | 7,8% | 30,8% |
| 49774 | 17,3% | 31,0% | 12,1% | 18,2% | 33,3% | 7,8% | 38,5% |
| 52703 | 14,8% | 20,7% | 3,0% | 36,4% | 6,7% | 11,8% | 0,0% |
| 51891 | 6,2% | 24,1% | 18,2% | 0,0% | 6,7% | 15,7% | 15,4% |
| 80423 | 35,8% | 65,5% | 54,5% | 36,4% | 70,0% | 68,6% | 69,2% |
| 78697 | 14,8% | 24,1% | 12,1% | 27,3% | 16,7% | 3,9% | 30,8% |
| 41583 | 16,0% | 20,7% | 12,1% | 27,3% | 13,3% | 9,8% | 23,1% |
| 124711 | 29,6% | 37,9% | 15,2% | 27,3% | 43,3% | 29,4% | 69,2% |

| |
|---|
| >50% |
| 20-49% |
| 1-19% |
| 0% |

**Suppl. Table 2.** Peptides identified in the differential serum binding pattern (DSBP) in the comparison between ELDERS and HCC patients with percentage of responders.

| UniqueID | Log2fc | pvalue | ELDERS | HCC | Species |
|---|---|---|---|---|---|
| pep_7853 | 3,04 | 0,0015288 | 45,28% | 23,53% | Rhino A |
| pep_21455 | 2,80 | 0,0083242 | 68,87% | 52,94% | Adeno C |
| pep_80349 | 2,79 | 0,0027759 | 83,96% | 84,31% | HHV-1 |
| pep_31975 | 2,79 | 0,0074886 | 57,55% | 33,33% | Rhino A |
| pep_30608 | 2,67 | 0,0020517 | 77,36% | 60,78% | HHV-1 |
| pep_55868 | 2,61 | 0,0043839 | 61,32% | 35,29% | HHV-4 |
| pep_30399 | 2,45 | 0,0026169 | 55,66% | 52,94% | HHV-1 |
| pep_37187 | 2,43 | 0,0049561 | 15,09% | 1,96% | Rota A |
| pep_21753 | 2,42 | 0,0068037 | 19,81% | 9,80% | Entero B |
| pep_95657 | 2,35 | 0,0087589 | 18,87% | 1,96% | Infl A |
| pep_56780 | 2,32 | 0,0025249 | 16,04% | 5,88% | HHV-5 |
| pep_41246 | 2,27 | 0,0025468 | 16,04% | 3,92% | Entero B |
| pep_56034 | 2,24 | 0,0093611 | 21,70% | 15,69% | HHV-4 |
| pep_6953 | 2,17 | 0,0090426 | 20,75% | 11,76% | HRSV |
| pep_60010 | 2,14 | 0,0071673 | 16,98% | 17,65% | HHV-4 |
| pep_58345 | 1,95 | 0,003944 | 16,98% | 1,96% | Adeno D |
| pep_80607 | 1,95 | 0,0006781 | 16,04% | 3,92% | Puumala virus |
| pep_47304 | 1,91 | 0,0022678 | 8,49% | 7,84% | HHV-6b |
| pep_55858 | 1,87 | 0,0039595 | 9,43% | 1,96% | HHV-4 |
| pep_15236 | 1,84 | 0,0061091 | 6,60% | 3,92% | HHV-1 |
| pep_56495 | 1,84 | 0,0073914 | 12,26% | 1,96% | HRSV |
| pep_70430 | 1,81 | 0,0029624 | 11,32% | 9,80% | HRSV |
| pep_70425 | 1,77 | 0,000883 | 14,15% | 7,84% | HRSV |
| pep_83385 | 1,77 | 0,0051046 | 14,15% | 0,00% | Infl B |
| pep_21385 | 1,66 | 0,00378 | 11,32% | 5,88% | Adeno C |
| pep_56411 | 1,65 | 0,0007978 | 16,98% | 1,96% | HRSV |
| pep_2940 | 1,58 | 0,0014241 | 13,21% | 5,88% | HRSV |
| pep_61544 | 1,49 | 0,0089181 | 10,38% | 1,96% | HHV-3 |
| pep_34232 | 1,49 | 0,0033276 | 13,21% | 1,96% | HHV-5 |
| pep_83955 | 1,49 | 0,0059948 | 11,32% | 1,96% | Infl A |
| pep_27705 | 1,43 | 0,0070204 | 12,26% | 1,96% | HIV-1 |
| pep_120234 | 1,32 | 0,0070106 | 9,43% | 0,00% | Borr burgd |
| pep_56859 | 1,32 | 0,006893 | 10,38% | 3,92% | HHV-5 |
| pep_111489 | 1,30 | 0,0028267 | 13,21% | 0,00% | Utive virus |
| pep_9115 | 1,26 | 0,0038686 | 8,49% | 0,00% | Adeno A |
| pep_52988 | 1,26 | 0,0075649 | 9,43% | 1,96% | HHV-4 |
| pep_93542 | 1,26 | 0,003347 | 9,43% | 1,96% | Rhino A |
| pep_110718 | 1,07 | 0,0091529 | 14,15% | 5,88% | Pegivirus A |
| pep_66028 | 1,00 | 0,003789 | 15,09% | 3,92% | MHV-1 |
| pep_54814 | 1,00 | 0,0083406 | 12,26% | 0,00% | PHV-2 |
| pep_88030 | -1,00 | 0,0046914 | 1,89% | 0,04% | HCV |
| pep_124703 | -1,06 | 0,0011823 | 1,89% | 3,92% | Staphyl aureus |
| pep_88950 | -1,24 | 0,0034665 | 0,94% | 0,00% | Entero A |
| pep_90150 | -1,35 | 0,0041204 | 1,89% | 0,00% | HEV |
| pep_85224 | -1,35 | 0,0022299 | 1,89% | 0,00% | Infl A |
| pep_89068 | -1,42 | 0,0022008 | 1,89% | 0,00% | Entero A |

| |
|---|
| >50% |
| 20-49% |
| 1-19% |
| 0% |

**Suppl. Table 3.** Peptides identified in the differential serum binding pattern (DSBP) in the comparison between ADULTS and THYR patients with percentage of responders.

| UniqueID | Log2fc | pvalue | ADULTS | THYROID | Species |
|---|---|---|---|---|---|
| pep_49774 | -5,77 | 1,60E-06 | 17,28% | 38,46% | Entero B |
| pep_87017 | -5,06 | 0,000489 | 20,99% | 38,46% | Entero B |
| pep_37254 | -4,99 | 0,0006587 | 17,28% | 23,08% | Entero B |
| pep_24523 | -4,94 | 0,0038019 | 45,68% | 53,85% | HHV-1 |
| pep_85871 | -4,82 | < 1e-07 | 11,11% | 30,77% | Entero B |
| pep_70404 | -4,69 | 0,0054304 | 44,44% | 53,85% | HHV-1 |
| pep_6954 | -4,62 | 0,0037516 | 24,69% | 30,77% | HRSV |
| pep_41583 | -4,61 | 5,90E-06 | 16,05% | 23,08% | HHV-6A |
| pep_37255 | -4,39 | 0,0014112 | 18,52% | 23,08% | Entero B |
| pep_43991 | -4,38 | 3,00E-07 | 2,47% | 30,77% | HPV 2 |
| pep_78697 | -4,36 | 0,0001812 | 14,81% | 30,77% | Entero B |
| pep_30426 | -4,14 | 0,0002766 | 9,88% | 15,38% | HHV-1 |
| pep_93248 | -4,09 | 9,41E-05 | 9,88% | 15,38% | INFL B |
| pep_124711 | -3,87 | 0,0088448 | 29,63% | 69,23% | STAPH AU |
| pep_58427 | -3,81 | < 1e-07 | 2,47% | 7,69% | Adenovirus F |
| pep_7028 | -3,81 | 0,0084402 | 1,23% | 0,00% | HRSV |
| pep_51732 | -3,80 | 0,0014067 | 6,17% | 0,00% | HHV-3 |
| pep_60340 | -3,65 | 4,10E-06 | 14,81% | 23,08% | HHV-4 |
| pep_81114 | -3,60 | 0,0017394 | 11,11% | 30,77% | INFL A |
| pep_121871 | -3,54 | 0,0057044 | 2,47% | 15,38% | Entamoeba histo |
| pep_21517 | -3,53 | 0,0030141 | 6,17% | 0,00% | Adenovirus C |
| pep_32104 | -3,47 | 0,0001627 | 11,11% | 23,08% | HHV-4 |
| pep_388 | -3,18 | 4,00E-07 | 8,64% | 23,08% | HHV-1 |
| pep_21036 | -3,15 | 0,0080248 | 4,94% | 7,69% | HHV-4 |
| pep_95377 | -3,06 | < 1e-07 | 7,41% | 23,08% | Entero C |
| pep_41650 | -3,00 | 0,0053844 | 3,70% | 23,08% | ROTA A |
| pep_32085 | -2,96 | 0,0003955 | 8,64% | 7,69% | HHV-4 |
| pep_12275 | -2,83 | 0,0002206 | 3,70% | 7,69% | HRSV |
| pep_115280 | -2,78 | 1,00E-07 | 8,64% | 30,77% | HHV-5 |
| pep_95657 | -2,76 | 7,00E-06 | 4,94% | 30,77% | INFL A |
| pep_25313 | -2,75 | 0,0001374 | 2,47% | 23,08% | INFL A |
| pep_124870 | -2,74 | 0,0028235 | 0,00% | 23,08% | INFL A |
| pep_3535 | -2,69 | 0,0004519 | 1,23% | 7,69% | INFL A |
| pep_5889 | -2,57 | 0,001088 | 4,94% | 0,00% | INFL A |
| pep_34174 | -2,54 | 0,007004 | 3,70% | 0,00% | INFL A |
| pep_13100 | -2,45 | < 1e-07 | 3,70% | 23,08% | INFL A |
| pep_21469 | -2,42 | 0,0005022 | 2,47% | 7,69% | INFL A |
| pep_121872 | -2,40 | 6,30E-06 | 2,47% | 23,08% | INFL A |
| pep_25542 | -2,38 | 0,0086106 | 2,47% | 0,00% | INFL A |
| pep_51891 | -2,37 | 0,0010971 | 6,17% | 15,38% | INFL A |
| pep_95294 | -2,37 | 6,20E-06 | 4,94% | 15,38% | INFL A |
| pep_14414 | -2,35 | 5,20E-06 | 1,23% | 7,69% | INFL A |
| pep_93699 | -2,35 | 7,60E-06 | 2,47% | 7,69% | INFL A |
| pep_52186 | -2,23 | 8,30E-05 | 0,00% | 7,69% | INFL A |
| pep_29978 | -2,22 | 5,40E-06 | 4,94% | 0,00% | INFL A |
| pep_59729 | -2,18 | 0,0011257 | 4,94% | 0,00% | INFL A |
| pep_25307 | -2,11 | 0,0002992 | 7,41% | 7,69% | INFL A |
| pep_72527 | -2,10 | 0,0002806 | 4,94% | 23,08% | INFL A |
| pep_64492 | -2,07 | 4,80E-06 | 0,00% | 15,38% | INFL A |
| pep_32721 | -2,04 | 0,0049091 | 0,00% | 7,69% | INFL A |
| pep_94230 | -2,00 | 0,0002634 | 2,47% | 30,77% | INFL A |
| pep_65097 | -2,00 | 0,0015626 | 0,00% | 15,38% | INFL A |
| pep_1707 | -1,97 | 0,001503 | 1,23% | 0,00% | INFL A |
| pep_56036 | -1,93 | 3,00E-07 | 0,00% | 7,69% | INFL A |

| |
|---|
| >50% |
| 20-49% |
| 1-19% |
| 0% |

| | | | | | |
|---|---|---|---|---|---|
| pep_6347 | -1,90 | 0,0002683 | 1,23% | 15,38% | INFL A |
| pep_112789 | -1,90 | 4,70E-06 | 1,23% | 15,38% | INFL A |
| pep_11958 | -1,86 | 0,0016081 | 1,23% | 0,00% | INFL A |
| pep_65643 | -1,86 | 0,0059359 | 1,23% | 0,00% | INFL A |
| pep_25092 | -1,85 | 0,0077421 | 0,00% | 7,69% | INFL A |
| pep_50487 | -1,81 | 0,0034373 | 0,00% | 15,38% | INFL A |
| pep_70425 | -1,81 | 0,0043427 | 2,47% | 15,38% | INFL A |
| pep_55557 | -1,77 | 0,0003338 | 1,23% | 7,69% | INFL A |
| pep_95097 | -1,75 | 0,0003177 | 1,23% | 7,69% | INFL A |
| pep_53237 | -1,75 | 0,0002873 | 1,23% | 23,08% | INFL A |
| pep_122146 | -1,72 | 0,0099109 | 0,00% | 7,69% | INFL A |
| pep_111489 | -1,72 | 0,0003129 | 0,00% | 15,38% | INFL A |
| pep_51029 | -1,68 | 0,0003109 | 1,23% | 0,00% | INFL A |
| pep_54814 | -1,68 | 0,000331 | 0,00% | 15,38% | INFL A |
| pep_128033 | -1,67 | 0,0069192 | 1,23% | 23,08% | INFL A |
| pep_52709 | -1,67 | 0,0092951 | 2,47% | 7,69% | INFL A |
| pep_68457 | -1,67 | 0,0005779 | 1,23% | 7,69% | INFL A |
| pep_56488 | -1,66 | 0,0002639 | 3,70% | 23,08% | INFL A |
| pep_71930 | -1,63 | 0,0002903 | 1,23% | 23,08% | INFL A |
| pep_22887 | -1,63 | 0,0002699 | 2,47% | 0,00% | INFL A |
| pep_6978 | -1,63 | 0,0002628 | 2,47% | 0,00% | INFL A |
| pep_70220 | -1,58 | 0,0002628 | 0,00% | 7,69% | INFL A |
| pep_115687 | -1,58 | 0,0002699 | 0,00% | 23,08% | INFL A |
| pep_125734 | -1,58 | 0,0002632 | 0,00% | 7,69% | INFL A |
| pep_61545 | -1,58 | 0,0002647 | 2,47% | 0,00% | INFL A |
| pep_11374 | -1,58 | 0,0002667 | 2,47% | 0,00% | INFL A |
| pep_93427 | -1,54 | 0,0002637 | 3,70% | 7,69% | INFL A |
| pep_121553 | -1,54 | 0,0002628 | 0,00% | 15,38% | INFL A |
| pep_81326 | -1,54 | 0,0002662 | 0,00% | 7,69% | INFL A |
| pep_40449 | -1,54 | 0,0002664 | 1,23% | 7,69% | INFL A |
| pep_59304 | -1,49 | 0,0002655 | 4,94% | 15,38% | INFL A |
| pep_43604 | -1,43 | 0,0002743 | 1,23% | 0,00% | HPV 11 |
| pep_63450 | -1,43 | 0,0002675 | 1,23% | 0,00% | HHV-5 |
| pep_16586 | -1,40 | 0,0002628 | 3,70% | 0,00% | RHINO B |
| pep_125400 | -1,40 | 0,0002629 | 1,23% | 0,00% | STREPT PNEU |
| pep_20964 | -1,38 | 0,0002637 | 1,23% | 7,69% | HHV-4 |
| pep_16322 | -1,38 | 1,60E-06 | 3,70% | 15,38% | HRSV |
| pep_63837 | -1,38 | 0,000489 | 3,70% | 0,00% | HRSV |
| pep_120923 | -1,32 | 0,0006587 | 0,00% | 7,69% | Candida albicans |
| pep_22876 | -1,32 | 0,0038019 | 0,00% | 7,69% | HHV-1 |
| pep_70403 | -1,32 | < 1e-07 | 1,23% | 0,00% | HHV-1 |
| pep_52988 | -1,32 | 0,0054304 | 0,00% | 7,69% | HHV-4 |
| pep_36724 | -1,32 | 0,0037516 | 0,00% | 15,38% | VACCINIA |
| pep_9238 | -1,26 | 5,90E-06 | 0,00% | 23,08% | HBV |
| pep_22259 | -1,26 | 0,0014112 | 1,23% | 7,69% | HRSV |
| pep_65949 | -1,26 | 3,00E-07 | 0,00% | 7,69% | MHV-1 |
| pep_124876 | -1,26 | 0,0001812 | 0,00% | 7,69% | STAPH AU |
| pep_3905 | -1,20 | 0,0002766 | 0,00% | 15,38% | Bat corona 1B |
| pep_30959 | -1,20 | 9,41E-05 | 0,00% | 7,69% | Adeno F |
| pep_77787 | -1,20 | 0,0088448 | 0,00% | 30,77% | HHV-6B |
| pep_47480 | -1,20 | < 1e-07 | 0,00% | 15,38% | SARS |
| pep_78570 | -1,20 | 0,0084402 | 1,23% | 30,77% | VEEV |
| pep_22694 | -1,14 | 0,0014067 | 1,23% | 46,15% | Adeno C |
| pep_18038 | -1,07 | 4,10E-06 | 0,00% | 15,38% | HPV 1 |
| pep_72905 | -1,07 | 0,0017394 | 0,00% | 30,77% | Moll CONT |
| pep_112256 | -1,00 | 0,0057044 | 0,00% | 7,69% | PEGI A |

**Suppl. Table 4.** Peptides identified in the differential serum binding pattern (DSBP) in the comparison between ELDERS and THOR patients with percentage of responders.

| UniqueID | Log2fc | pvalue | ELDERS | THO | Species |
|---|---|---|---|---|---|
| pep_52703 | -4,18 | 0,0001 | 6,60% | 20,69% | HHV- 4 |
| pep_57549 | -3,66 | 0,0002 | 10,38% | 17,24% | HHV- 4 |
| pep_58457 | -3,63 | 0,0002 | 21,70% | 48,28% | ADENO A |
| pep_2496 | -3,56 | 0,0021 | 63,21% | 58,62% | HRSV |
| pep_41583 | -3,51 | 0,0016 | 10,38% | 20,69% | HHV- 6A |
| pep_23621 | -3,48 | 0,0000 | 6,60% | 27,59% | RHINO A |
| pep_2510 | -3,43 | 0,0001 | 20,75% | 13,79% | HRSV |
| pep_95378 | -3,32 | 0,0021 | 13,21% | 24,14% | Entero C |
| pep_79241 | -2,96 | 0,0026 | 11,32% | 31,03% | Entero B |
| pep_7048 | -2,95 | 0,0055 | 31,13% | 41,38% | HRSV |
| pep_95805 | -2,78 | 0,0002 | 2,83% | 10,34% | INFL A virus |
| pep_47472 | -2,78 | 0,0036 | 17,92% | 13,79% | SARS |
| pep_32085 | -2,66 | 0,0012 | 5,66% | 20,69% | HHV- 4 |
| pep_23579 | -2,51 | 0,0003 | 6,60% | 17,24% | RHINO A |
| pep_20858 | -2,49 | 0,0036 | 13,21% | 20,69% | HHV- 4 |
| pep_843 | -2,48 | 0,0001 | 6,60% | 10,34% | RHINO B |
| pep_26629 | -2,36 | 0,0003 | 3,77% | 0,00% | HHV- 3 |
| pep_41220 | -2,13 | 0,0006 | 2,83% | 10,34% | HHV- 4 |
| pep_8005 | -2,13 | 0,0000 | 2,83% | 10,34% | ADENOB |
| pep_72402 | -2,09 | 0,0001 | 3,77% | 6,90% | HHV- 8 |
| pep_95259 | -1,97 | 0,0021 | 4,72% | 10,34% | INFL A virus |
| pep_58735 | -1,97 | 0,0019 | 2,83% | 13,79% | Entero B |
| pep_81682 | -1,97 | 0,0022 | 1,89% | 3,45% | INFL B virus |
| pep_53023 | -1,97 | 0,0001 | 0,94% | 3,45% | HHV- 4 |
| pep_47676 | -1,90 | 0,0001 | 2,83% | 6,90% | HHV- 4 |
| pep_114671 | -1,87 | 0,0006 | 2,83% | 6,90% | MERS |
| pep_6978 | -1,84 | 0,0008 | 5,66% | 3,45% | HRSV |
| pep_30857 | -1,79 | 0,0001 | 1,89% | 3,45% | HHV- 1 |
| pep_87124 | -1,74 | 0,0009 | 2,83% | 3,45% | Encephalom virus |
| pep_35810 | -1,70 | 0,0030 | 2,83% | 0,00% | INFL A virus |
| pep_20726 | -1,69 | 0,0034 | 4,72% | 0,00% | HHV- 4 |
| pep_95329 | -1,63 | 0,0001 | 2,83% | 3,45% | Entero C |
| pep_33713 | -1,63 | 0,0001 | 0,94% | 0,00% | HHV- 5 |
| pep_49775 | -1,58 | 0,0057 | 3,77% | 17,24% | Entero B |
| pep_25836 | -1,58 | 0,0099 | 1,89% | 6,90% | HHV- 3 |
| pep_38647 | -1,54 | 0,0024 | 2,83% | 10,34% | ADENOC |
| pep_72879 | -1,54 | 0,0013 | 0,94% | 0,00% | Molluscum contagiosum |
| pep_54192 | -1,54 | 0,0012 | 0,94% | 6,90% | Papiine herpesvirus 2 |
| pep_81079 | -1,50 | 0,0034 | 3,77% | 0,00% | INFL A virus |
| pep_18809 | -1,49 | 0,0007 | 1,89% | 10,34% | Vaccinia virus |
| pep_61334 | -1,42 | 0,0057 | 4,72% | 3,45% | Betapapillomavirus 1 |
| pep_90150 | -1,40 | 0,0021 | 1,89% | 13,79% | Hepatitis E virus |
| pep_51051 | -1,35 | 0,0014 | 2,83% | 0,00% | Rabies virus |
| pep_56866 | -1,30 | 0,0032 | 25,47% | 24,14% | HHV- 5 |
| pep_21656 | -1,30 | 0,0007 | 2,83% | 27,59% | RHINO B |
| pep_10619 | -1,27 | 0,0086 | 0,94% | 6,90% | Hepatitis E virus |
| pep_17535 | -1,27 | 0,0097 | 0,94% | 17,24% | Mamastrovirus 1 |
| pep_37286 | -1,24 | 0,0074 | 3,77% | 3,45% | Entero B |
| pep_95507 | -1,20 | 0,0007 | 0,94% | 0,00% | Entero C |
| pep_125342 | -1,20 | 0,0007 | 0,00% | 3,45% | Streptococcus pneu |
| pep_96000 | -1,17 | 0,0100 | 6,60% | 3,45% | INFL A virus |
| pep_92766 | -1,14 | 0,0008 | 0,94% | 0,00% | INFL A virus |
| pep_24988 | -1,13 | 0,0037 | 2,83% | 0,00% | HPV 9 |
| pep_32056 | -1,13 | 0,0074 | 1,89% | 0,00% | HHV- 4 |

| |
|---|
| >50% |
| 20-49% |
| 1-19% |
| 0% |

| | | | | | |
|---|---|---|---|---|---|
| pep_47865 | -1,06 | 0,0083 | 0,94% | 0,00% | Vaccinia virus |
| pep_17407 | -1,06 | 0,0099 | 2,83% | 0,00% | Zaire ebolavirus |
| pep_44839 | -1,00 | 0,0091 | 1,89% | 0,00% | ADENOF |
| pep_47070 | -1,00 | 0,0071 | 1,89% | 0,00% | HHV- 7 |
| pep_78467 | -1,00 | 0,0056 | 2,83% | 3,45% | INFL A virus |

**Suppl. Table 5.** Peptides identified in the differential serum binding pattern (DSBP) in the comparison between ADULTS and THOR patients with percentage of responders.

| UniqueID | Log2fc | pvalue | ADULTS | THOR | SPECIES |
|---|---|---|---|---|---|
| pep_41583 | -5,66178 | 1,00E-07 | 16,05% | 20,69% | HHV- 6A |
| pep_52703 | -5,12102 | 5E-07 | 14,81% | 20,69% | HHV- 4 |
| pep_58457 | -4,56294 | 9E-07 | 11,11% | 48,28% | ADENOA |
| pep_2510 | -4,49185 | 1,00E-07 | 4,94% | 13,79% | HRSV |
| pep_124711 | -4,05889 | 0,00027 | 29,63% | 37,93% | STAPHYL AU |
| pep_55868 | -4,04182 | 0,00095 | 41,98% | 55,17% | HHV- 4 |
| pep_57549 | -3,926 | 7,15E-05 | 16,05% | 17,24% | HHV- 4 |
| pep_70408 | -3,91709 | 0,000375 | 24,69% | 44,83% | HHV- 1 |
| pep_58754 | -3,90689 | 0,000314 | 20,99% | 31,03% | Entero B |
| pep_95378 | -3,76445 | 0,000546 | 17,28% | 24,14% | Entero C |
| pep_23621 | -3,7625 | 1,3E-06 | 8,64% | 27,59% | RHINO A |
| pep_78697 | -3,63005 | 0,000067 | 14,81% | 24,14% | Entero B |
| pep_95330 | -3,62149 | 0,00255 | 38,27% | 31,03% | Entero C |
| pep_16674 | -3,58986 | 0,001213 | 22,22% | 55,17% | HRSV |
| pep_51891 | -3,54432 | 1,8E-06 | 6,17% | 24,14% | HRSV |
| pep_52914 | -3,43296 | 0,001632 | 22,22% | 41,38% | HHV- 4 |
| pep_17601 | -3,365 | 0,000966 | 17,28% | 44,83% | HRSV |
| pep_85617 | -3,35509 | 0,001808 | 20,99% | 31,03% | Entero B |
| pep_25315 | -3,30505 | 0,005021 | 28,40% | 48,28% | HHV- 5 |
| pep_52926 | -3,22239 | 0,008185 | 32,10% | 48,28% | HHV- 4 |
| pep_47473 | -3,10434 | 0,005409 | 17,28% | 20,69% | SARS |
| pep_94986 | -3,07682 | 0,002439 | 20,99% | 24,14% | Entero C |
| pep_47472 | -3,04891 | 0,00207 | 13,58% | 13,79% | SARS |
| pep_81422 | -3,03242 | 0,000672 | 11,11% | 17,24% | INFL A virus |
| pep_57555 | -3,02237 | 0,0003 | 6,17% | 13,79% | HHV- 4 |
| pep_60340 | -3 | 0,004065 | 14,81% | 27,59% | HHV- 4 |
| pep_81351 | -3 | 0,000103 | 7,41% | 13,79% | INFL A virus |
| pep_122510 | -2,98089 | 0,000515 | 8,64% | 31,03% | Helicobacter pylorI |
| pep_11867 | -2,96963 | 8,5E-06 | 6,17% | 10,34% | HRSV |
| pep_56479 | -2,94642 | 0,006376 | 22,22% | 51,72% | HRSV |
| pep_25382 | -2,90689 | 0,000233 | 7,41% | 34,48% | HHV- 1 |
| pep_7840 | -2,848 | 0,002527 | 9,88% | 20,69% | RHINO A |
| pep_8572 | -2,80735 | 0,001546 | 11,11% | 10,34% | Hepatitis B virus |
| pep_24481 | -2,79008 | 3,81E-05 | 2,47% | 17,24% | HHV- 1 |
| pep_115288 | -2,7625 | 0,000121 | 7,41% | 10,34% | HHV-5 |
| pep_32085 | -2,73697 | 0,001716 | 8,64% | 20,69% | HHV- 4 |
| pep_21036 | -2,73697 | 0,000831 | 4,94% | 13,79% | HHV- 4 |
| pep_94735 | -2,72247 | 5E-07 | 2,47% | 10,34% | INFL A virus |
| pep_843 | -2,70044 | 4,73E-05 | 4,94% | 10,34% | RHINO B |
| pep_115278 | -2,69302 | 0,000362 | 6,17% | 13,79% | HHV-5 |
| pep_730 | -2,65535 | 2,3E-06 | 2,47% | 3,45% | HRSV |
| pep_121870 | -2,64386 | 0,008802 | 11,11% | 24,14% | Entamoeba histolytica |
| pep_83385 | -2,58496 | 9,1E-06 | 1,23% | 3,45% | INFL B virus |
| pep_72527 | -2,54057 | 5,07E-05 | 4,94% | 17,24% | HHV- 8 |
| pep_41933 | -2,53343 | 0,002313 | 9,88% | 10,34% | Human parainfl 1 |
| pep_81114 | -2,53343 | 0,00253 | 11,11% | 27,59% | INFL A virus |
| pep_93542 | -2,51457 | 0,000661 | 3,70% | 10,34% | RHINO A |
| pep_20858 | -2,48543 | 0,006771 | 9,88% | 20,69% | HHV- 4 |
| pep_94230 | -2,48543 | 3,5E-06 | 2,47% | 20,69% | INFL A virus |
| pep_6953 | -2,47393 | 0,00842 | 11,11% | 17,24% | HRSV |
| pep_26629 | -2,46949 | 0,000783 | 2,47% | 0,00% | HHV- 3 |
| pep_60009 | -2,45943 | 0,002006 | 9,88% | 20,69% | HHV- 4 |
| pep_95805 | -2,45943 | 0,003938 | 8,64% | 10,34% | INFL A virus |
| pep_23579 | -2,42033 | 0,002478 | 6,17% | 17,24% | RHINO A |

| Legend |
|---|
| >50% |
| 20-49% |
| 1-19% |
| 0% |

| | | | | | |
|---|---|---|---|---|---|
| pep_37187 | -2,38827 | 0,003237 | 7,41% | 13,79% | Rotavirus A |
| pep_41220 | -2,37346 | 0,000147 | 3,70% | 10,34% | HHV- 4 |
| pep_124875 | -2,36923 | 0,000226 | 2,47% | 0,00% | STAPHYL AU |
| pep_115280 | -2,35755 | 0,00277 | 8,64% | 20,69% | HHV-5 |
| pep_56780 | -2,3505 | 4,8E-06 | 2,47% | 27,59% | HHV- 5 |
| pep_95259 | -2,32193 | 0,000158 | 1,23% | 10,34% | INFL A virus |
| pep_93556 | -2,29546 | 1,41E-05 | 4,94% | 13,79% | RHINO A |
| pep_93995 | -2,24101 | 0,000206 | 3,70% | 13,79% | INFL A virus |
| pep_63822 | -2,22239 | 0,000608 | 3,70% | 6,90% | HRSV |
| pep_58914 | -2,21501 | 0,001362 | 4,94% | 27,59% | Entero B |
| pep_24228 | -2,18763 | 0,00685 | 16,05% | 27,59% | Entero C |
| pep_32245 | -2,18442 | 0,009078 | 8,64% | 24,14% | HHV- 5 |
| pep_731 | -2,16993 | 0,001068 | 4,94% | 10,34% | HRSV |
| pep_81151 | -2,152 | 0,008112 | 8,64% | 24,14% | INFL A virus |
| pep_30762 | -2,14975 | 0,006948 | 7,41% | 10,34% | HHV- 1 |
| pep_37950 | -2,14684 | 0,002229 | 6,17% | 13,79% | Entero C |
| pep_92942 | -2,14296 | 0,000242 | 4,94% | 10,34% | INFL A virus |
| pep_25316 | -2,1375 | 0,009982 | 6,17% | 41,38% | HHV- 5 |
| pep_19082 | -2,12928 | 0,004389 | 7,41% | 31,03% | Sindbis virus |
| pep_82552 | -2,11548 | 0,000305 | 1,23% | 6,90% | Rabies virus |
| pep_120590 | -2,10692 | 0,001445 | 3,70% | 3,45% | Bos taurus (Bovine) |
| pep_58427 | -2,10434 | 6,3E-06 | 2,47% | 17,24% | ADENOF |
| pep_11376 | -2,08746 | 0,00096 | 2,47% | 24,14% | HHV- 5 |
| pep_118117 | -2,07039 | 3E-07 | 0,00% | 6,90% | Pegivirus A |
| pep_83949 | -2,03242 | 0,000294 | 1,23% | 13,79% | INFL A virus |
| pep_85871 | -2 | 0,001058 | 11,11% | 27,59% | Entero B |
| pep_81342 | -2 | 0,000172 | 1,23% | 6,90% | INFL A virus |
| pep_58735 | -1,97199 | 0,003792 | 4,94% | 13,79% | Entero B |
| pep_6978 | -1,96683 | 0,000843 | 2,47% | 3,45% | HRSV |
| pep_121869 | -1,96347 | 3,06E-05 | 1,23% | 3,45% | Entamoeba histolytica |
| pep_14414 | -1,96347 | 4,4E-06 | 1,23% | 13,79% | Orf virus |
| pep_94108 | -1,926 | 0,000118 | 1,23% | 0,00% | INFL A virus |
| pep_81697 | -1,91427 | 0,003993 | 4,94% | 13,79% | INFL B virus |
| pep_25073 | -1,90046 | 0,007384 | 7,41% | 31,03% | RHINO A |
| pep_57910 | -1,89812 | 0,000997 | 3,70% | 17,24% | RHINO A |
| pep_5222 | -1,88982 | 0,009761 | 4,94% | 6,90% | ADENOC |
| pep_22259 | -1,88753 | 1,77E-05 | 1,23% | 6,90% | HRSV |
| pep_60278 | -1,848 | 0,008888 | 7,41% | 24,14% | HHV- 5 |
| pep_53023 | -1,8413 | 0,003486 | 3,70% | 3,45% | HHV- 4 |
| pep_95377 | -1,80735 | 0,00459 | 7,41% | 10,34% | Entero C |
| pep_28530 | -1,80735 | 0,000116 | 1,23% | 10,34% | Coronavirus HKU1 |
| pep_20726 | -1,80735 | 0,003556 | 4,94% | 0,00% | HHV- 4 |
| pep_34175 | -1,79355 | 0,009109 | 3,70% | 3,45% | HHV- 5 |
| pep_47676 | -1,77259 | 0,004834 | 3,70% | 6,90% | HHV- 4 |
| pep_22243 | -1,77259 | 0,002562 | 2,47% | 17,24% | HRSV |
| pep_22694 | -1,76553 | 2,53E-05 | 1,23% | 10,34% | ADENOC |
| pep_30959 | -1,76553 | 2,3E-06 | 0,00% | 13,79% | ADENOF |
| pep_29959 | -1,76553 | 2,93E-05 | 0,00% | 10,34% | HHV- 4 |
| pep_95657 | -1,75899 | 0,006129 | 4,94% | 27,59% | INFL A virus |
| pep_120706 | -1,72247 | 0,00055 | 1,23% | 6,90% | Bos taurus (Bovine) |
| pep_81573 | -1,72247 | 3,32E-05 | 2,47% | 17,24% | INFL B virus |
| pep_53143 | -1,70044 | 0,004212 | 4,94% | 10,34% | HHV- 4 |
| pep_16322 | -1,67807 | 0,000132 | 3,70% | 10,34% | HRSV |
| pep_72879 | -1,67807 | 0,000542 | 1,23% | 0,00% | Molluscum contagiosum |
| pep_125606 | -1,67807 | 0,000153 | 1,23% | 0,00% | Toxoplasma gondii |
| pep_49755 | -1,66985 | 0,00089 | 4,94% | 27,59% | Entero B |
| pep_61545 | -1,66985 | 0,001412 | 2,47% | 6,90% | HHV- 3 |

| | | | | | |
|---|---|---|---|---|---|
| pep_43991 | -1,65711 | 0,00523 | 2,47% | 37,93% | HPV 2 |
| pep_25542 | -1,63227 | 0,000142 | 2,47% | 10,34% | HHV- 1 |
| pep_54814 | -1,63227 | 2,67E-05 | 0,00% | 13,79% | Papiine herpesvirus 2 |
| pep_18809 | -1,63227 | 0,000608 | 3,70% | 10,34% | Vaccinia virus |
| pep_63836 | -1,62803 | 0,001504 | 3,70% | 6,90% | HRSV |
| pep_81079 | -1,62803 | 0,00236 | 2,47% | 0,00% | INFL A virus |
| pep_91466 | -1,58496 | 0,000555 | 0,00% | 3,45% | INFL A virus |
| pep_41650 | -1,58496 | 0,001398 | 3,70% | 13,79% | Rotavirus A |
| pep_61334 | -1,54057 | 0,002416 | 1,23% | 3,45% | HPV 1 |
| pep_32215 | -1,54057 | 0,000806 | 2,47% | 34,48% | HHV- 5 |
| pep_82064 | -1,54057 | 0,00357 | 3,70% | 17,24% | Human parainfl 3 |
| pep_54192 | -1,54057 | 0,004352 | 4,94% | 6,90% | Papiine herpesvirus 2 |
| pep_25337 | -1,53605 | 0,000198 | 1,23% | 3,45% | HHV- 1 |
| pep_55991 | -1,53605 | 0,000135 | 0,00% | 3,45% | HHV- 4 |
| pep_17599 | -1,53605 | 0,000182 | 0,00% | 6,90% | HRSV |
| pep_71674 | -1,53605 | 0,000135 | 0,00% | 3,45% | Orf virus |
| pep_23067 | -1,53605 | 0,00057 | 1,23% | 6,90% | Rotavirus A |
| pep_61544 | -1,48543 | 0,00012 | 0,00% | 3,45% | HHV- 3 |
| pep_18185 | -1,48543 | 9,62E-05 | 0,00% | 3,45% | HHV- 8 |
| pep_111578 | -1,48543 | 0,000108 | 0,00% | 10,34% | Salivirus A |
| pep_34273 | -1,44746 | 0,001085 | 1,23% | 20,69% | HHV- 5 |
| pep_46163 | -1,44746 | 0,005322 | 3,70% | 6,90% | HHV- 7 |
| pep_65058 | -1,44746 | 0,007683 | 1,23% | 3,45% | HHV- 8 |
| pep_96000 | -1,43296 | 0,000654 | 1,23% | 3,45% | INFL A virus |
| pep_52585 | -1,39855 | 0,001467 | 4,94% | 13,79% | Coronavirus HKU1 |
| pep_52977 | -1,39855 | 0,007991 | 2,47% | 0,00% | HHV- 4 |
| pep_53891 | -1,39855 | 0,004673 | 1,23% | 3,45% | Orf virus |
| pep_16554 | -1,39855 | 0,006346 | 3,70% | 20,69% | RHINO B |
| pep_37286 | -1,37851 | 0,000749 | 0,00% | 3,45% | Entero B |
| pep_32298 | -1,37851 | 0,000663 | 0,00% | 10,34% | HHV- 2 |
| pep_40449 | -1,37851 | 0,000839 | 1,23% | 10,34% | JEV |
| pep_124094 | -1,37851 | 0,000825 | 0,00% | 6,90% | Porphyr gingivalis |
| pep_56959 | -1,36923 | 0,00561 | 1,23% | 6,90% | HRSV |
| pep_109939 | -1,34792 | 0,005107 | 3,70% | 0,00% | MERS |
| pep_51051 | -1,34792 | 0,008729 | 3,70% | 0,00% | Rabies virus |
| pep_2940 | -1,32193 | 0,006192 | 1,23% | 17,24% | HRSV |
| pep_36042 | -1,32193 | 0,000667 | 0,00% | 10,34% | ADENOA |
| pep_34232 | -1,32193 | 0,000102 | 0,00% | 13,79% | HHV- 5 |
| pep_46641 | -1,32193 | 0,000795 | 1,23% | 3,45% | HHV- 6A |
| pep_65047 | -1,32193 | 0,000667 | 0,00% | 3,45% | HHV- 8 |
| pep_83964 | -1,32193 | 0,003103 | 1,23% | 3,45% | INFL A virus |
| pep_93121 | -1,32193 | 0,003072 | 1,23% | 10,34% | INFL A virus |
| pep_32450 | -1,32193 | 0,000601 | 0,00% | 10,34% | INFL B virus |
| pep_81699 | -1,32193 | 0,000662 | 1,23% | 3,45% | INFL B virus |
| pep_60927 | -1,32193 | 0,00071 | 0,00% | 3,45% | INFL C virus |
| pep_65097 | -1,29546 | 0,000915 | 0,00% | 17,24% | Molluscum contagiosum |
| pep_24988 | -1,26303 | 0,003191 | 1,23% | 0,00% | HPV 9 |
| pep_89064 | -1,26303 | 0,003086 | 1,23% | 0,00% | Entero A |
| pep_67160 | -1,26303 | 0,000645 | 0,00% | 3,45% | HPV 3 |
| pep_50960 | -1,26303 | 0,003083 | 0,00% | 20,69% | ADENO D |
| pep_9388 | -1,26303 | 0,003133 | 0,00% | 10,34% | HHV- 5 |
| pep_57926 | -1,26303 | 0,003095 | 0,00% | 10,34% | RHINO A |
| pep_93596 | -1,26303 | 0,00315 | 1,23% | 20,69% | RHINO A |
| pep_18827 | -1,26303 | 0,003095 | 1,23% | 0,00% | Vaccinia virus |
| pep_60516 | -1,24101 | 0,009194 | 4,94% | 0,00% | HHV- 6B |
| pep_95040 | -1,20163 | 0,003138 | 1,23% | 0,00% | Entero C |
| pep_10824 | -1,20163 | 0,000632 | 0,00% | 10,34% | ADENO A |

| | | | | | |
|---|---|---|---|---|---|
| pep_54291 | -1,20163 | 0,000758 | 0,00% | 13,79% | ADENOB |
| pep_25894 | -1,20163 | 0,003076 | 1,23% | 3,45% | HHV- 3 |
| pep_83398 | -1,20163 | 0,003237 | 0,00% | 13,79% | INFL B virus |
| pep_17391 | -1,20163 | 0,003169 | 1,23% | 0,00% | Orf virus |
| pep_115284 | -1,18442 | 0,007423 | 1,23% | 17,24% | HHV-5 |
| pep_16814 | -1,1375 | 0,0001 | 0,00% | 17,24% | ADENOE |
| pep_34625 | -1,1375 | 0,003277 | 0,00% | 6,90% | HHV- 5 |
| pep_7124 | -1,1375 | 0,003395 | 1,23% | 0,00% | HHV- 8 |
| pep_63837 | -1,1375 | 0,003109 | 3,70% | 13,79% | HRSV |
| pep_84917 | -1,1375 | 0,000561 | 0,00% | 3,45% | INFL A virus |
| pep_53986 | -1,1375 | 0,000859 | 0,00% | 10,34% | Orf virus |
| pep_115113 | -1,1375 | 0,000874 | 0,00% | 20,69% | Pegivirus A |
| pep_82438 | -1,12553 | 0,008402 | 4,94% | 17,24% | Measles virus |
| pep_15376 | -1,07039 | 0,00313 | 2,47% | 17,24% | ADENOF |
| pep_65049 | -1,07039 | 0,003862 | 0,00% | 3,45% | HHV- 8 |
| pep_108859 | -1,07039 | 0,003463 | 1,23% | 3,45% | MERS |
| pep_110718 | -1,07039 | 0,000151 | 0,00% | 17,24% | Pegivirus A |
| pep_125760 | -1,07039 | 0,000548 | 0,00% | 6,90% | Toxoplasma gondii |
| pep_78570 | -1,07039 | 0,004287 | 1,23% | 6,90% | VEEV |
| pep_25296 | -1,06413 | 0,008029 | 1,23% | 13,79% | HHV- 5 |
| pep_121780 | -1 | 0,00361 | 0,00% | 3,45% | Ehrlichia chaffeensis |
| pep_8944 | -1 | 0,000613 | 0,00% | 10,34% | HHV- 8 |
| pep_7028 | -1 | 0,003333 | 1,23% | 10,34% | HRSV |
| pep_81712 | -1 | 0,003396 | 0,00% | 3,45% | INFL B virus |
| pep_21715 | -1 | 0,004213 | 2,47% | 0,00% | RHINO B |

**Suppl. Table 6.** Peptides identified in the differential serum binding pattern (DSBP) in the comparison between ELDERS and SARC patients with percentage of responders.

| UniqueID | Log2fc | pvalue | ELDERS | SARC | Species |
|---|---|---|---|---|---|
| pep_93748 | 4,38 | 0,000112 | 44,34% | 27,27% | RHINO A |
| pep_93426 | 4,08 | 0,000167 | 65,09% | 51,52% | RHINO A |
| pep_6954 | 3,42 | 0,002301 | 41,51% | 24,24% | HRSV |
| pep_20958 | 3,26 | 0,00537 | 30,19% | 27,27% | HHV-4 |
| pep_58992 | 3,10 | 0,008094 | 29,25% | 30,30% | Entero B |
| pep_85617 | 3,10 | 0,005894 | 16,04% | 12,12% | Entero B |
| pep_21029 | 2,96 | 0,009554 | 48,11% | 54,55% | HHV-4 |
| pep_31975 | 2,94 | 0,004008 | 57,55% | 48,48% | RHINO A |
| pep_21753 | 2,88 | 0,007115 | 19,81% | 15,15% | Entero B |
| pep_95657 | 2,87 | 0,001354 | 18,87% | 6,06% | INFL A virus |
| pep_12301 | 2,69 | 0,004321 | 11,32% | 6,06% | HHV-4 |
| pep_25047 | 2,66 | 0,002514 | 16,98% | 6,06% | RHINO A |
| pep_41933 | 2,40 | 0,006046 | 10,38% | 6,06% | PARAINFL virus 1 |
| pep_51737 | 2,10 | 0,00397 | 14,15% | 3,03% | HHV-3 |
| pep_58427 | 2,07 | 0,005013 | 15,09% | 9,09% | ADENOF |
| pep_22741 | 2,00 | 0,006585 | 41,51% | 30,30% | HHV-1 |
| pep_83385 | 1,77 | 0,007257 | 14,15% | 6,06% | INFL B virus |
| pep_111489 | 1,45 | 0,00777 | 13,21% | 0,00% | Utive virus |
| pep_86085 | -1,32 | 0,000259 | 1,89% | 3,03% | Dengue virus |
| pep_42655 | -1,40 | 0,000254 | 1,89% | 0,00% | Entero C |
| pep_81079 | -1,47 | 0,002543 | 3,77% | 3,03% | INFL A virus |

>50%
20-49%
1-19%
0%

**Suppl. Table 7.** Peptides identified in the differential serum binding pattern (DSBP) in the comparison between ADULTS and SARCOMA patients with percentage of responders.

| UniqueID | Log2fc | pvalue | ADULTS | SARC | Species |
|---|---|---|---|---|---|
| pep_93426 | 3,37 | 0,003623 | 59,26% | 51,52% | RHINO A |
| pep_93748 | 3,32 | 0,006482 | 51,85% | 27,27% | RHINO A |
| pep_88487 | -1,00 | 0,005723 | 2,47% | 0,00% | HCV |
| pep_54290 | -1,00 | 0,006602 | 1,23% | 3,03% | ADENOB |
| pep_31000 | -1,00 | 0,005973 | 0,00% | 6,06% | INFL B virus |
| pep_73800 | -1,00 | 0,005709 | 2,47% | 0,00% | Molluscum contagiosum |
| pep_20897 | -1,14 | 0,00567 | 3,70% | 3,03% | HHV-4 |
| pep_7028 | -1,26 | 0,001395 | 1,23% | 9,09% | HRSV |
| pep_41650 | -1,30 | 0,006481 | 3,70% | 0,00% | Rotavirus A |
| pep_11867 | -1,46 | 0,008159 | 6,17% | 12,12% | HRSV |
| pep_92942 | -1,46 | 0,006627 | 4,94% | 3,03% | INFL A virus |
| pep_81079 | -1,54 | 0,001634 | 2,47% | 3,03% | INFL A virus |
| pep_94230 | -1,58 | 0,000283 | 2,47% | 3,03% | INFL A virus |
| pep_94735 | -1,58 | 0,000295 | 2,47% | 0,00% | INFL A virus |
| pep_25542 | -1,63 | 0,000319 | 2,47% | 12,12% | HHV-1 |
| pep_25307 | -1,66 | 0,008419 | 7,41% | 9,09% | HHV-5 |
| pep_81437 | -1,72 | 0,000283 | 2,47% | 0,00% | INFL A virus |
| pep_82552 | -1,77 | 0,002005 | 1,23% | 0,00% | Rabies virus |
| pep_12275 | -1,85 | 0,000264 | 3,70% | 12,12% | HRSV |
| pep_125606 | -1,96 | 5,93E-05 | 1,23% | 6,06% | Toxoplasma gondii |

>50%
20-49%
1-19%
0%
